# Actin contractility and endocytosis create apico-basal tension gradient in HeLa cells

**DOI:** 10.1101/2025.02.17.638627

**Authors:** Tanmoy Ghosh, Anisha Majhi, Avijit Kundu, Ayan Banerjee, Bidisha Sinha

## Abstract

The functions of the plasma membrane of a cell are coupled to its mechanical state. To understand the players that contribute to the tight regulation of membrane mechanics with its local heterogeneities, the tension of the cell membrane requires to be mapped out from the basal to the apical membrane. In this work, optical-trap-based tension measurements performed at two axial (z) planes per cell reveal an apico-basal gradient, that is understood better on comparison with basal membrane mechanics and membrane compaction measurements by a combination of high resolution microscopy (IRM) and fluorescence lifetime (Flipper-TR) measurements, respectively. While the apico-basal gradient in the apparent tension (low to high) was found to depend on an intact actin cytoskeleton, analysing the colocalization of the cortical actin with the motor protein myosin and the actin-membrane linker ezrin suggested that cytoskeleton contractility is a major determinant of the patterned tension. Tension-sensitive lifetime of Flipper-TR, validated the gradient but failed to catch the dependence on actin due to the dual and opposite effects of tension on lifetime. Finally, measurements utilizing fluorescently tagged transferrin (Tf) demonstrated that the functional state of the membrane also showed similar height dependence. Thus, planes where tension maximized showed a peak in accumulation of Tf colocalizing with clathrin indicative of the presence of more pits. This enabled us to demonstrate that cells exist in tightly regulated patterned tension states that is coupled to their cytoskeleton as well membrane functions like endocytosis.

**Significance Statement:** The cell membrane is protective but also highly dynamic - performing a diverse set of functions usually via membrane shape changes. These functions affect membrane mechanics but are also mechano-regulated. Mapping the state of cell surface mechanics or tension of live cells is therefore of fundamental importance. Interestingly, recent works have suggested that membrane tension can be non-uniform. However, no single tension-measuring technique can truly reach every part of the cell. Our work addresses this issue where we map tension in live HeLa cells using multifarious approaches including the measurement of tether forces, membrane fluctuations, and lipid compaction. We show that at the top/apical surface, the membrane-tension is lower than that at the midplane. The observed gradient resembles that of the state of endocytosis and controlled acto-myosin contractility.

## Introduction

Tension at the cell surface is a crucial parameter in cell biophysics because it plays a fundamental role in various cellular processes and functions ^1–5^. Membrane tension influences cell shape and structure, cell motility and migration ^6^, membrane trafficking ^7^, cell signalling, endo/exocytosis, mechanosensation and the overall cellular homeostasis ^8–10^. Understanding the mechanical properties of cells, including tension, is critical for insights into biophysical mechanisms underlying functional cellular processes as well as mechanisms responsible for malignancies^11^. Despite the extensive body of work demonstrating the role of tension in various functional contexts, very few studies focus on intracellular distribution of tension which can organize processes in cells^12^. Although tension has been traditionally believed to equilibrate instantaneously, multiple recent reports point to the possibility of slow equilibration of tension in cells ^13–15^. With the tension being measured by extracting nanoscale tethers from cells using optically trapped polystyrene beads, these reports show how while due to certain functional requirements some cells prefer faster equilibration of tension ^16^, other cells may have extremely slow rates of tension equilibration. Slow equilibration raises the possibility of inhomogeneities – or, the maintenance of distinct tension states at different regions in single cells. Whether such states exist – if they are random stable variations or are they are patterned – are some questions that remain unanswered due to the technical limitations of measuring tension at multiple points in single cells.

Various techniques can be used to measure cell membrane tension. Optical trap-based tether pulling assay^17,18^, Atomic Force Microscopy (AFM)^19,20^ and imaging-based methods ^21,22^ can be utilized for estimating membrane tension ^23,24^. Optical tweezers^25^ use a focused laser beam to trap and manipulate small objects, such as microbeads, attached to the cell membrane. These beads serve as handles to apply and measure forces and reveal the membrane tension from the tether force. However, it includes an extra cost of breaking membrane cytoskeleton bonds, therefore tension is expected to be higher than the basal in-plane membrane tension. Hence, the tension obtained is termed as the ‘apparent’ membrane tension. Due to the geometry of the experiment, measuring tension at multiple locations in same cells in order to map out tension, is challenging and induces phototoxicity in cells if conducted at many locations. In addition, OT-based tension measurements cannot be performed at the base of adherent cells since tethers cannot be drawn there – thereby setting a limit to the reach of this technique. On the other hand, interference reflection microscopy (IRM) can map the "effective" tension of the basal membrane with a xy resolution set by a typical diffraction-limited image from the nanoscopic z-fluctuations it measures of the membrane^22,26^. However, IRM measures spontaneous membrane fluctuations (including active fluctuations in addition to thermal fluctuations^27^) to derive the effective fluctuation tension and cannot to applied to the lateral/apical cell membrane. In contrast to both these techniques, fluorescence-based tension reporters (Flipper-TR) can reach any part of the cell^28^. However, the interpretation of tension from the fluorescence lifetime is non-trivial and sometimes cannot be deconvolved from the effect of any altered lipid order. Thus, to gauge the level of heterogeneities and patterns that exist in the membrane mechanics axially along a cell, a combination of tools may become necessary.

Any heterogeneity or pattern in membrane mechanics in cells would need active participation of various machinery. The actin-cytoskeleton has been shown to be critical in the flow of tension^29^. In the cortical cytoskeleton underlying the plasma membrane, the non-muscle myosin II serve as the molecular motors that walk on actin as a bundled fibre and thereby contract the actin network^30^ through dynamic binding and power strokes. The effect of this contractility on membrane tension has been studied and reported to both enhance or reduce tension in a context dependent manner^8^. On the other hand, the actin-membrane coupler protein – Ezrin – has been shown to be required to maintain a higher tensional state^19^. In addition to actin, the endocytic state of the membrane is also intertwined with its mechanical state^4,31^. Recent work validates the fact that making new clathrin coated pits (CCPs) from free fluctuating membrane reduces the excess area stored in the microscopic membrane undulations – thus increasing tension^32^ while reports in polarized cells reveal region-specific control of endocytosis^33,34^.

In this work, we have combined OT-based tether extraction, IRM-based measurement of fluctuations tension, and Flipper-TR-based lipid compaction measurements to demonstrate the basal and apico-basal patterns in tension in single cells. Utilizing confocal and STED microscopy we propose that actin contractility and coated pits are major determinants of this apico-basal tension gradient, while membrane actin attachment by ezrin plays a weaker role. Multiple physical measurements point to midplane to apical plane decrease in tension, and also suggest that the functional state of the cell tightly couples with its mechanics since regions of higher tension reveal more accumulation of CCPs in comparison to internalized/clustered cargo.

## Results

### Coupling optical trap to IRM shows similar estimation of cell surface tension across a population

In order to have a better overall assessment of tension heterogeneities, we first coupled two distinct measurement techniques on the same imaging platform (**Fig. 1A**). This integration enabled the sequential measurement of two different types of tension at different positions within the same cell. To have consistent cell shapes and to have a well-presented lateral surface away from base, cells were grown of line-shaped micropatterns (width – 20 µm, spacing – 15 µm). The optical trap (OT) based tether extractions were performed at the sides of cells (lateral cell surface) using trapped polystyrene microspheres (**Fig. 1B left, Fig, S1**). Using IRM, we measured the membrane fluctuations of the basal membrane of the same cell, therefore reporting the effective fluctuation tension ^35^ of the basal surface. During the tether extraction, events of rupture (of possibly a multi-tether system) were also noticed (cyan arrow, phase 4, **Fig. 1B centre**). In such cases, the force jump was considered as the tether force^18^. To confirm that the final plateau force corresponded to force for a single tether, the extraction velocity was increased at the final stage of the measurement. This caused the tether to break releasing the bead from the tether such that it moved back to the centre of the trap (red arrow, phase 5, **Fig. 1B centre**). The final rupture, resulting in zero force, confirmed that the last force plateau (100-200 sec, **Fig. 1B centre**) was indeed due to a single tether, which confirmed that, single-tether forces were accurately measured in these experiments. Additionally, the z-position of the trapped bead was also recorded for each extraction. By subtracting the z-position of the substrate (top of the coverslip), the relative z-distance from the substrate at which the tether was extracted could be estimated (“Height”, **Fig. 1C**).

**Figure 1:**
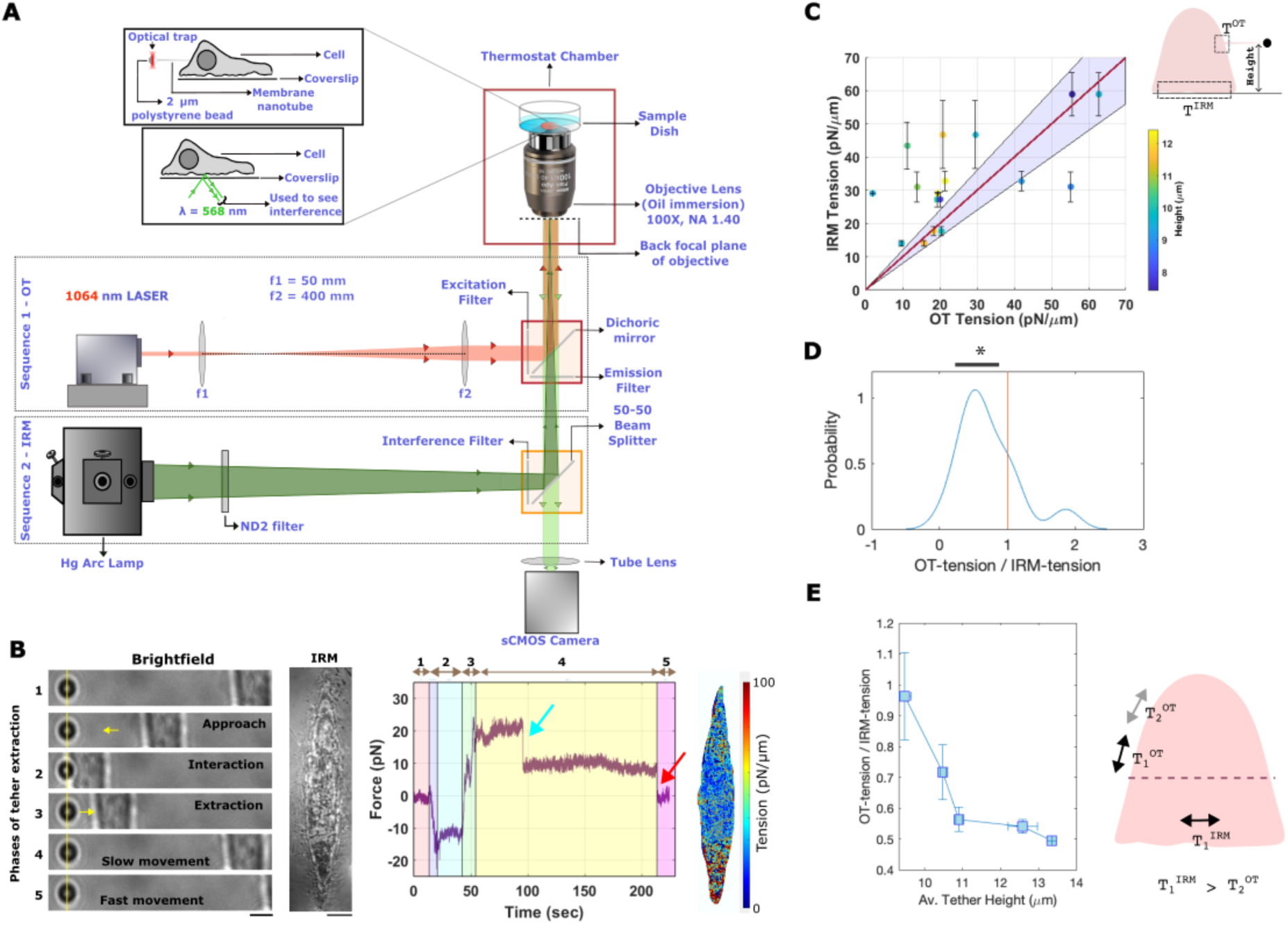
Optical trap and IRM based apparent tension measurements in HeLa cells. A) Schematic representation of the ray diagram illustrating sequential imaging setup for Total Internal Reflection Fluorescence (TIRF) and Interference Reflection Microscopy (IRM). B) Left (first) panel: Images of a typical cell undergoing sequential steps of manipulation for tether extraction. Second panel: Bead shown is trapped by the optical tweezer. Yellow line is a guide to the eye to show bead centre. IRM image of the same cells manipulated in the left. Third (central) panel: Plot of force felt by the bead at different phases of tether extraction and indicated by numbers at the top that correspond to image number of left panel. Fourth (right) panel: Map of tension of the basal membrane obtained of IRM image series of same cell shown in first-to-third panel. S*calebar* – 5 μm. C) Comparison of basal membrane tension (IRM tension) and apical membrane tension (OT tension). The color bar indicates the height of the tether from the cell’s basal surface. The red line represents the y = x reference line, while the violet-shaded region denotes a 20% deviation in the slope from this line. Number of independent repeats = 3. N_cells_= 7. Adjacent schematic dep “height” of tether D) Distribution of ratio of tension (measured by OT and tension measured by IRM on same single cells). * denotes the ratio is statistically different than 1 with a p-value <0.05. E) Dependence of ratio of tension (measured by OT and tension measured by IRM on same single cells) on the average tether height (of the OT measurements on that cell). Right schematic depicts the regions assessed by IRM and OT and that T obtained by IRM is usually higher than the T obtained from OT from very apical regions of the cell.

Following the tether extraction, the basal membrane fluctuations of the same cells were imaged at 20 fps (2048 frames). Temporal variations in IRM intensity (ΔI) at the same pixel locations were converted to membrane height fluctuations (Δh) using previously established methodology^22,26^. These nanometric membrane height fluctuations were then used to estimate the effective fluctuation tension at the basal surface (**Fig. 1B right**). We next compared the median fluctuation tension at the basal plane and the apparent tension at the equatorial planes, which are located at higher height than basal membrane for multiple single cells. A comparison of OT-tension and IRM-Tension (**Fig. 1C**) revealed that both techniques yielded tension values of the same order of magnitude. However, no significant correlation could be captured, likely due to the distinct nature of the two measurement techniques. The comparison of the average (median) amplitude of temporal fluctuations (SD_time_, **Fig. S2**) for each cell versus the showed a negatively correlated (with Spearman correlation coefficient of -0.3) but non-significant trend.

Despite the lack of correlation, we examined the distribution of the ratio of OT tension to IRM tension measured for every cell individually. For most cells OT tension was lower than IRM tension, with the ratio typically being less than one (**Fig. 1D**). We next checked if the tether height could also influence the OT-based tension measurement. Although colour-coding the tether height (**Fig. 1 C**) did not reveal any clear trend, averaging the data using a sliding window along the x-axis (height) revealed a distinct height dependence. Specifically, the ratio of OT apparent membrane tension to the basal membrane IRM tension decreased as the tether height increased beyond 9 µm, indicating a height-dependent variation in OT-derived apparent tension (**Fig. 1 E**). It is important to note that the IRM tension at the basal membrane itself exhibited significant variability (**Fig. 1B right, Fig. S3**). While pixel-wise mapping of tension (**Fig. 1B right**) and other fit parameters (**Fig. S3**) was used for visualizations, for comparison between and IRM and OT measurements the median tension obtained from specific regions (FBRs, see Methods) was used.

To further investigate the height dependence of OT tension, we next measured tension at multiple locations on the lateral surface within the same cells.

### The lateral cell surface has a lower effective tension away from the substrate

In order to assess the state of tension at multiple positions in the same cell, two tethers were extracted per cell. Following a successful tether extraction, a separate bead was trapped to extract a second tether at a different height from the substrate (referred to as H_1_ and H_2_ in the schematic – **Fig. 2A**). For these measurements, the heights varied between 5 – 13 mm from the substrate (please put the correct values). As shown in **Fig. 2A**, the beads were positioned at a considerable distance from each other, avoiding regions near the nucleus. Sequential tether extraction yielded two tether forces per cell (**Fig. 2B**, lines connect measurements from same cells, Height 1 < Height 2). A negative correlation was observed between the tether force and its height (Pearson correlation coefficient= -0.5, p value= 0.009), indicating consistent decrease in tether force with increasing height (**Fig. 2C**). On the average, the tether force decreased by approximately ∼ 1.2 ± 0.5 pN/µm as tether height increased, as determined from single-cell comparisons across 54 cells from more than three independent experiments. The ratio of forces between lower and higher height tethers was found to be significantly higher from 1 for control cells (**Fig. 2D**). The fractional change per µm was found be ∼ 0.13 ± 0.11 indicating around 13% change per µm (**Fig. 2E**). Due to the cell-to-cell heterogeneity, experiments involving a single tether per cell were insufficient to detect the correlation between tether force and height. Therefore, sequential dual-tether experiments were essential for uncovering the apico-basal gradient in apparent tension.

**Figure 2:**
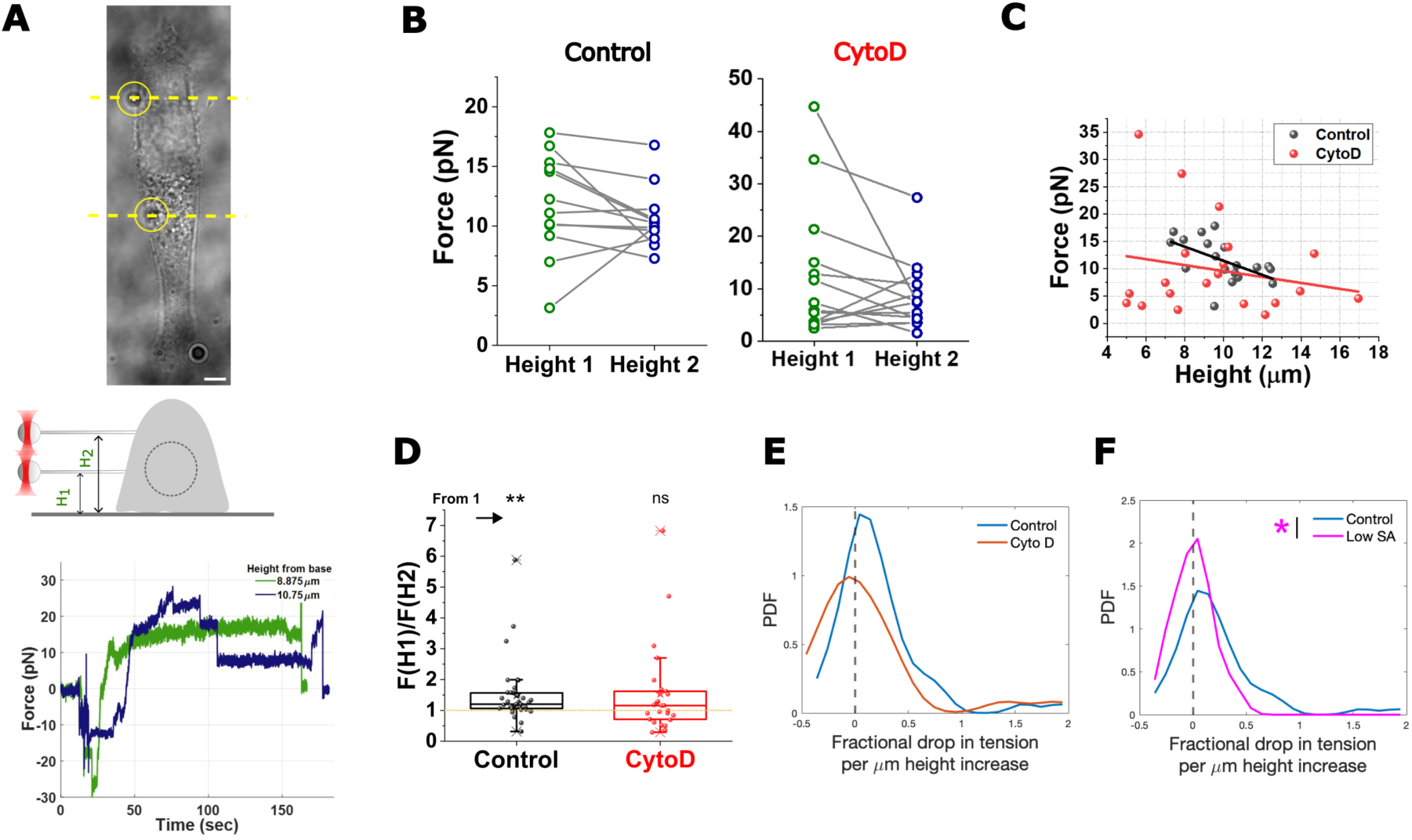
Measured apparent tension reduces at higher heights in an actin-dependent manner. A) Top: Representative brightfield image of cell with beads attached. Yellow circles denote the two beads used for the tether extractions. Dotted line are imaginary lines across which cross-sections may be drawn to provide the schematic in the middle. Middle: Schematic shows that the two beads were used to draw out tethers at two different heights. Bottom: typical force curves from same cell at two different heights performed sequentially. B) Plot of forces measured at two different heights for control and Cyto D treated cells from a representative set (cell numbers: N_control_ = 12, N_Cyto D_ = 14). Lines connect measurements from same cells. No. of independent repeats = 5 for each condition. C) Dependence of tether force on height for control and Cyto D treated cells. (N_Control_ = 12 cells; N_CytoD_ = 14 cells from 3 independent sets). D) Comparison of ratio forces from the lower height to the higher height. Arrow denotes comparison to test if ratios are different than 1. Ns represents non-significant different statistical difference between Control and Cyto D ratios. Data taken from 5 independent sets. (N_Control_ = 37 cells; N_Cyto D_ = 25 cells). Mann-Whitney test was performed and ** used to denote p-value < 0.01 and ns used to denote p-value>0.05. E, F) Probability distribution function (pdf) of fractional drop in tension per µm compared between E) Control and Cyto D-treated cells and F) Control and cells with low spread area (Low SA). Pink * denote significant difference with p-value<0.05 obtained using Mann-Whitney test. No. of independent sets = 5 (Control, Cyto D) and 2 sets (Low SA) data, Number of cells: N_Control_ = 37 cells; N_CytoD_ = 25 cells; N_low SA_ = 13.

### Actin-dependent correlation of effective tension with distance of the tether from the basal plane

The cortical actomyosin cytoskeleton is essential for maintaining tensional homeostasis in cells as it anchors the plasma membrane to the underlying contractile actin network. The configuration of this network influences cellular tension and may exhibit an apico-basal gradient in actin thickness as previously reported in CHO. Using 3D confocal scanning, we confirmed the presence of a similar gradient in HeLa cells. To further investigate the dependence of the tension gradient or force drop on the cortical actin, we treated cells with Cyto D, which disassembles the cortical actin network and performed tension measurements. Following Cyto D treatment, the correlation between the tether force and height was diminished (**Fig. 2 B, C**), as indicated by a reduction in the correlation coefficient statistically non-significant relation (Pearson correlation coefficient= -0.17, p value= 0.39). The average force drop per µm increase in height decreased to ∼ 0.2 ± 0.8 pN/µm, where – in this case, we varied the heights where beads were tethered to the cell even further from the substrate – between 5-17 µm. Additionally, the ratio of forces (**Fig. 2 D**) and the fractional tension change per µm increase in height (**Fig. 2E**) were not significantly different than one and zero respectively. However, the correlation and the fractional tension change became more heterogenous, potentially disrupting the smoothness of the gradient. In contrast, sequential dual-tether experiments performed in cells with lower spread area (**Fig. 2 F, low SA** (**Fig.S4**)), revealed a clearer and less noisy reduction in fractional tension change per µm. Together this section demonstrates that while actin reinforces the gradient, minimizing noise, cell attachment and spreading on the substrate also play an important role in establishing the observed tension gradient.

To further explore how actin differentially influences tension at varying heights, we next investigated the gradients in myosin-dependent cytoskeleton contractility and ezrin-mediated actin-membrane interactions.

### Stronger apico-basal gradient of actin’s colocalization with myosin than p-ezrin

To capture the gradient in contractility and the gradient in membrane-cortex attachment, F-actin was labelled with phalloidin while myosin or p-ezrin were labelled using immunofluorescence. 3D imaging across z-planes was performed (**Fig. 3 A**) to assess their distribution and colocalization. Cyto D was used to enhance depolymerization and induce network breakage (**Fig. 3 A bottom**). Line-patterned cells were fixed, and imaged using confocal microscopy equipped with 2D STED-based super-resolution. To evaluate the overlap of myosin and actin intensity, which marks regions of contractile actin, we quantified the fraction of the cortical actin network (∼ pixels within 400 nm of cell periphery) that contained myosin. Cells were imaged with a lateral pixel-size of 40 nm and a z-step of 100 nm. The lateral cortical actin was extracted and object-detection algorithms were applied to calculate the percentage of overlap of intense regions of actin with myosin/p-ezrin (**Fig. 3 B**). Endogenous non-muscle myosin (**Fig. 3 C**) and p-ezrin (**Fig. 3 D**) were separately assayed.

**Figure 3:**
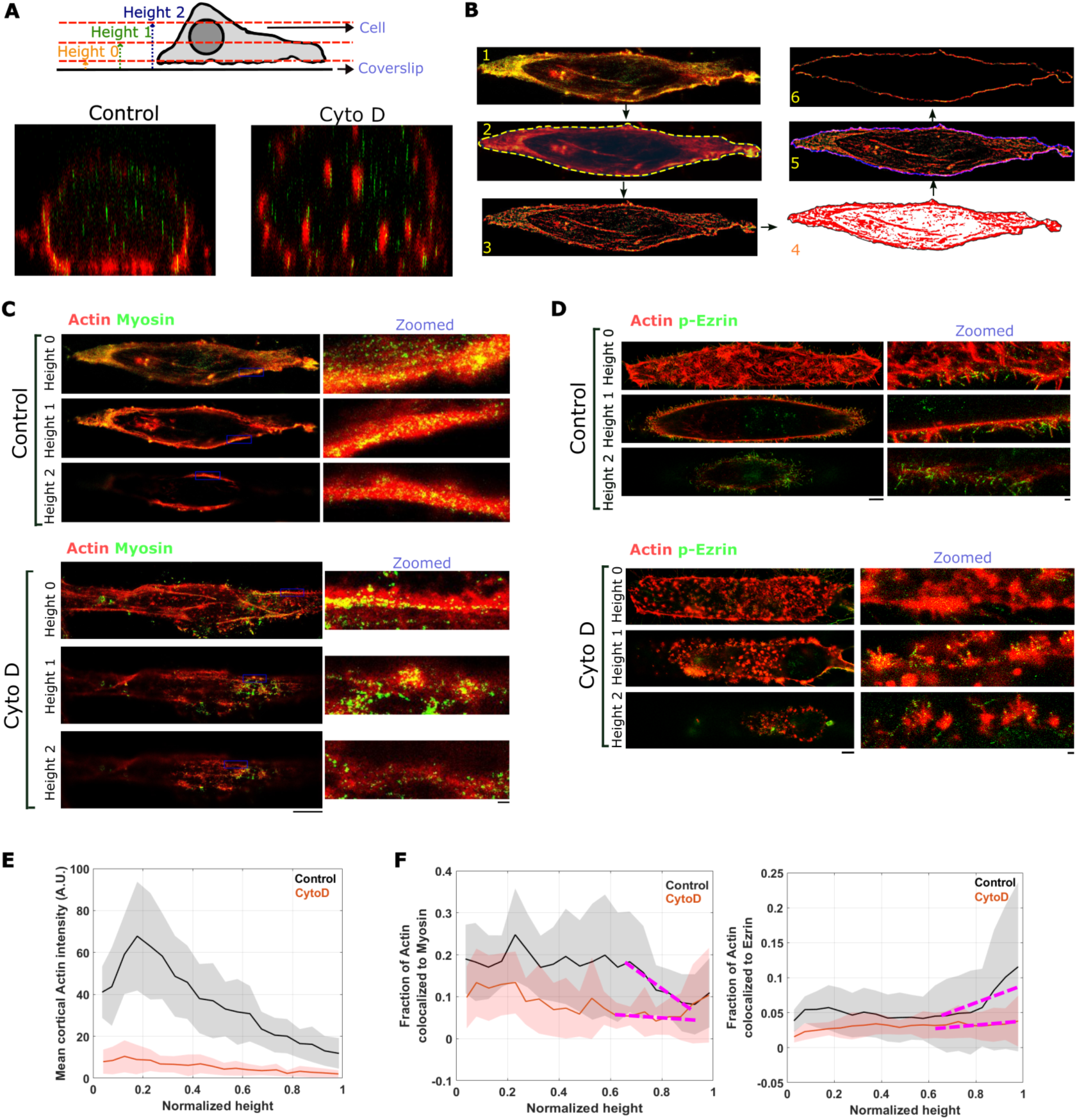
Enhanced colocalization of actin and myosin observed at lower planes. A) Top: Representative schematic of a cell’s cross section and the different imaging planes at different heights that will be used in subsequent sections of the figure. Bottom: X-Z projection of STED images of the actin (Alexa fluor 568 phalloidin) a control and a Cyto D-treated cell at three different heights. B) Depiction of the procedure of analysis of colocalization between F -actin and myosin/p-ezrin at regions within 40 nm of cell’s periphery. (1) Raw image used for the analysis of the colocalization. Red represents actin, green represents the myosin and yellow represents the colocalization. (2) An ROI (yellow dashed line) was drawn cell to select the cell. (3) Represents the binary images used for object detection, which was done by global thresholding. (4) A more accurate ROI was drawn along the boundary of the cell by taking the object as actin. (5) An ROI of 10 pixels (1 pix = 40 nm) thickness (shown in magenta) was made to collect the colocalization data near the cell’s membrane or the cortex region. (6) ROI area in (5) is shown in the image and this image is finally taken for the colocalization analysis. C, D) Representative images (with zoomed sections in the right) of planes at different heights on Control and Cyto D-treated conditions depicting F-actin (green) and C) non-muscle myosin II A (red) or D) p-ezrin. The yellow regions represent colocalization. Height 0 represents the plane at the base of the cell, Height 1 is an apical region above the base and Height 2 is much above the base. The cartoon represents heights relative to the coverslip of the adherent cell. *Unzoomed scalebar* – 5 μm, *Zoomed scalebar* – 0.5 μm. E) Height profile of mean actin intensity at the colocalized region at the cortex. Height is normalized such that base is 0 and top is 1. Shaded region denote SEM. Black: control; Red: Cyto D. F) Height profile of fraction of colocalization of actin to myosin (left) or p-Ezrin (right) at the cortex. Black line represents control and the red line represents the Cyto D treated cell. Height is normalized such that base is 0 and top is 1. Shaded region denote SEM. Black: control; Red: Cyto D. Data represents imaging from single set done for 10 control and 9 Cyto D treated cells. Dashed pink lines represent guides to eye pointing out the trend. For left (myosin) plot: Control: correlation coefficient: -0.34 (p val:0.0022); Cyto D: correlation coefficient: -0.1(p val:0.438)); for right plot: Control: correlation coefficient: 0.1954 (p val:0.0041); Cyto D: correlation coefficient: 0.0039 (p val:0.96)).

Interestingly, the cortical actin intensity increased as one moved away from the base, before showing a clear decline (**Fig. 3 E**). It is important to note that the height of the cell has been normalized to assess the gradients. Now, it is well known that fixation followed by mounting processes can lead to the shrinkage of cell heights. Thus, to avoid direct comparison to absolute heights of tethers pulled out of live cells, we present the dependence of intensity and colocalization on normalized heights. Cyto D resulted in a reduction in the cortical actin intensity as expected – weakening the absolute intensity differences along height (**Fig. 3 E**). In addition visualizing the overlap of myosin or p-Ezrin with actin in zoomed-in regions (**Fig. 3C, D**) could reveal the higher overlap for myosin than p-ezrin and the reduction of colocalized areas with height. Quantification of the colocalization of cortical actin with myosin revealed that typically 20% of the actin overlapped with myosin in lower planes (**Fig. 3 F left**). In contrast, only 5% of actin colocalized with p-ezrin (**Fig. 3 F right**). At higher planes, the overlap reduced to ∼ 10% (pink dashed line, **Fig. 3 F left** depicts the gradient (Pearson correlation coefficient: -0.34, P value: 0.002)). P-Ezrin staining did not show significant reduction, rather, the overlap increased with height (**Fig. 3 F right;** (Pearson correlation coefficient: 0.2, P value: 0.004)). The concentration (mean intensity) of actin in overlapping acto-myosin puncta at the cortex was observed to peak at 20% of the height of the cell and then reduced at higher planes. The gradients for acto-myosin colocalization fraction as well as actin-p-ezrin fraction were lowered after Cyto D treatment and the correlation coefficients were found to be non-significant (**Table S1**).

Myosin to actin ratio reflects the contractility of the network^30^. Therefore, it can be concluded from this section that contractility has a strong apico-basal gradient and could contribute more to the tension gradient than membrane-actin attachment.

### Apico-basal gradient in lipid compaction

We next investigated whether these forces also correlate with lipid compaction measured by the fluorescence lifetime of Flipper-TR, an indirect reporter of cell tension (**Fig. 4 A, B**). The lifetime of Flipper-TR can be influenced by changes in the lipid environment, such as alterations in axial rotation, which may become more or less restrictive (**Fig. 4 B**). As previously reported^28^, while increased tension could directly enhance lipid-lipid distances thereby lowering the lifetime, an opposing effect – tension-induced lipid ordering – can also reduce the average lifetime (**Fig. 4 B right**). Therefore, the observed changes in lifetime may not be linearly proportional to the change in tension – but influenced by a combination of factors. However, within single cells, the effect of enhancing tension has generally been observed to increase the lifetime, as the impact of altered lipid order tends to override the direct effect of tension.

**Figure 4:**
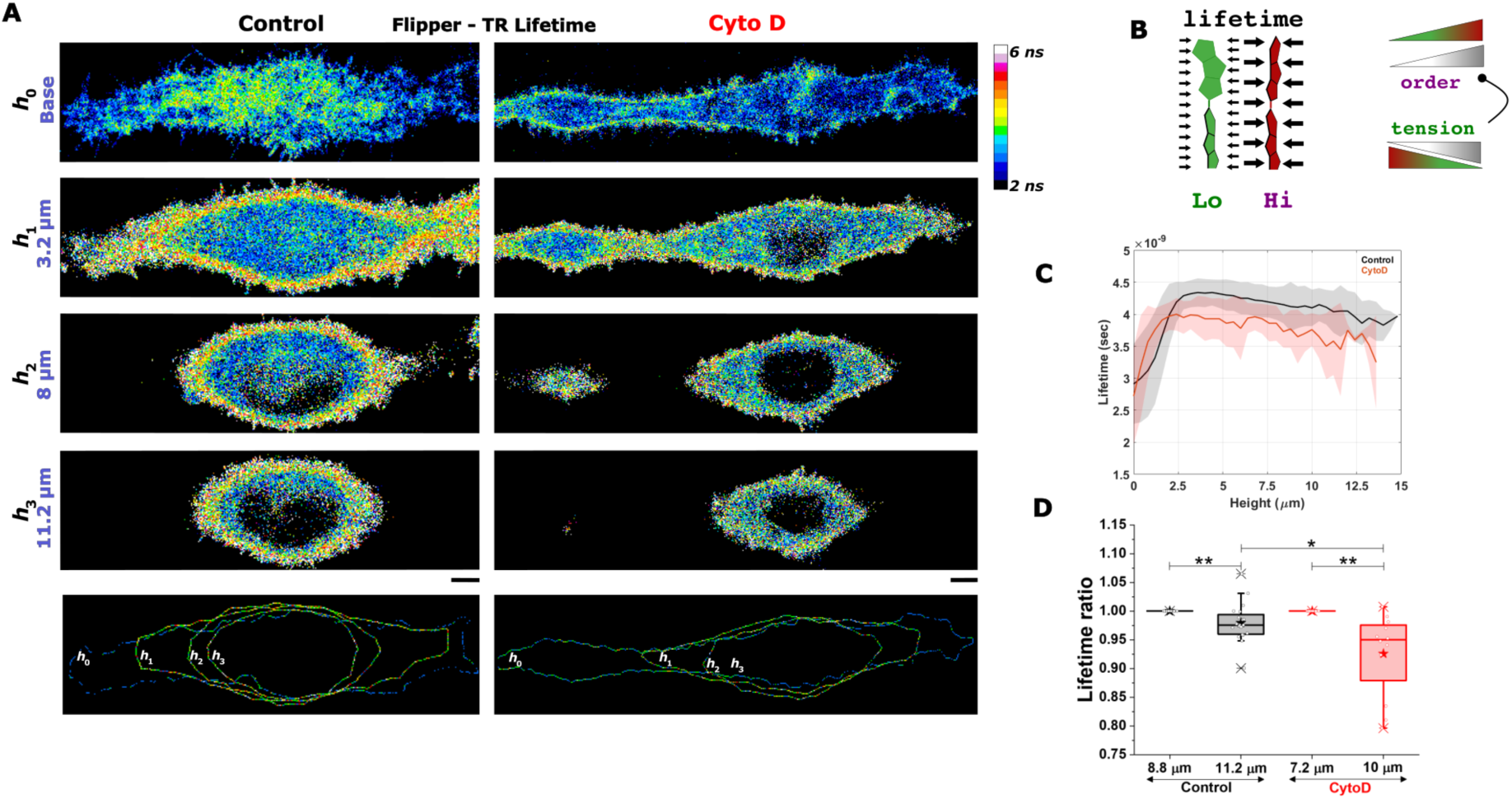
Apicobasal lifetime profile of Flip-TR confirms the tension gradient. A) Representative images of fluorescence lifetime of Flipper-TR (at different imaging planes) in line-micropatterned live HeLa cells. B) Left: Schematic showing how higher lateral pressure inhibits independent movement of the rotators of Flipper-TR increasing lifetime; Right: Schematic showing how increasing tension or order (gray scale gradients) alter the fluorescence lifetime (green (low lifetime) to maroon (high lifetime) of Flip-TR. Enhancing tension enhances order – as captured by the dot-headed arrow. C) Height-dependence of fluorescence lifetime of Flipper-TR for Control and Cyto D-treated cells. Solid line depicts the median trend while shaded region denotes the SEM. N=20 cells and 12 cells for control and Cyto D respectively. D) Significance check of the lifetime ratio between mid and higher imaging planes for Control and Cyto D-treated cells. Heights used are average heights used for the OT-based tension measurement for each case. Mann-Whitney test was performed and ** used to denote p-value < 0.01 and ns used to denote p-value>0.05.

Using 3D imaging of lifetime of Flipper-TR with confocal microscopy and a time-correlated single-photon counting module, we observed an initial increase in lifetime, followed by a decline as the imaging plane moved higher in the cell. Interestingly, the height at which the lifetime reached its peak closely matched the peak in cortical actin intensity (**Fig 3E**). At the base of the cell, the lifetime was significantly lower compared to higher planes (**Fig. 4 A, C**) and with increased cell-to-cell variation (SEM across cells, error bars in **Fig. 4 C**). For the analysis, only the peripheral membrane region (200 nm from edge) from all z-planes were extracted (**Fig. 4 A last row**). A weak but statistically significant apico-basal gradient in lifetime was observed, with higher planes showing significantly lower lifetime. In addition to the complete profile, the averages at z-planes corresponding to the average heights used in OT experiments were also compared. Normalization of height was not done since both OT and Flipper-TR used live cell experiments. However, for each cell, the change in lifetime with height was noted, and the lifetime ratio determined for comparison.

In line with OT-based measurements (**Fig. 2C**), Cyto D treated cells have lower lifetime akin to lower tension (**Fig. S5A**) and also display higher cell-to-cell variability (**Fig. S5B**). The lowering of lifetime with increasing height at the apical section was observed to be significant (**Fig. 4 D)**. Averaged across 34 cells, we found ∼ 1% reduction in lifetime per µm – reflecting the gradient in tension. In Cyto D-treated cells, however, this reduction increased to 1.7% per µm.

Tension gradient, measured by OT, reduced on Cyto D-treatment. To understand why there was no reduction (rather an increase) in the lifetime gradient in the Flipper-TR data (**Fig. 4D**), we have to take into account the opposing effects (of reducing tension) on lifetime - especially because Cyto D treatment lowers the basal tension. We believe that in Cyto D treated cells, the direct effect of lowering tension on lipid-lipid distance becomes less pronounced than in Control cells, because lipid-lipid distances are already minimized. Thus, at low basal tension, lifetime changes induced by order are more pronounced such that a lower tension gradient may show up as a higher lifetime-gradient

### Apico-basal gradient in Transferrin-loaded clathrin-coated endocytic pits

Both OT-based measurements and lifetime of Flipper-TR indicate a drop in tension as we proceed from the midplane towards the apical plane. However, Flipper-TR fails to reflect the loss of gradient on Cyto D treatment. Although the dual role of tension in affecting fluorescence lifetime – either directly or through lipid order – could explain the results, to further validate the mid-apical tension drop, we next checked its functional footprint. Endocytosis is known to affect tension as well as be influenced by tension ^36^. Here, we chose to map clathrin mediated endocytosis (CME) across the difference z-planes of the cells. To do so, a fluorescently labelled cargo of CME – Transferrin (Tf) was administered to the cells for 5 min. Subsequently cells were fixed and the endogenous clathrin was labelled by immunofluorescence. 3D STED imaging ensured a proper demarcation of the small puncta in not only the xy plane, but also in z. It must be noted that in CCPs, the cargo and clathrin are expected to colocalize or be in very close proximity. Once internalized, the clathrin coat dissociates from the Tf - loaded vesicles. Thus, we expect that if CCPs are higher than internalized cargo OT just-clustered cargo, the percentage Tf-rich regions colocalizing with clathrin would increase.

The first level visualization of the Tf puncta at different z done by z-dependent colour coding (**Fig. S6**) show that Tf-puncta at the midplane (yellow) appear bigger, while in Cyto D treated cells puncta appear smaller overall. To properly capture the trends, object-detection followed by image analysis was performed. On analysing the z-profile of Transferrin-rich regions (termed Tf objects/puncta), clathrin-rich pixels (termed Cl objects/puncta) and their colocalization (**Fig. S7, S8, Table S2**) in control, and Cyto D-treated cells (**Fig. 5A, B, Fig. S9**), two points were found to be robust. Firstly, the number density of Tf puncta (**Fig. 5C**) and its area fraction (**Fig. S9 A**) was highest at the midplane while reducing at lower and higher planes. clathrin also showed a similar profile (**Fig. S9 B,C**). Not only did Tf and Cl individually have higher densities at midplanes (reduced minimum distance between nearest neighbours - **Fig. S9 D, E**)), the colocalized area fraction also peaked at midplanes and reduced at higher apical surfaces (**Fig. 5D**). This implied that midplanes must have relatively more Tf-loaded clathrin coated pits contributing to the enhanced tension compared to higher planes. Note that while ∼20-30% Tf colocalized with clathrin (**Fig. S9F**), only 3-4% of the clathrin puncta colocalized with Tf (**Fig. S9G**). The closeness of nearest Tf and clathrin puncta also shows an increase in the midplanes (decreased minimum distance) followed by reduction at the apical planes (**Fig. S9H**). On Cyto D treatment, the number density dropped at all planes and there was no z-dependence in either of the parameters. Now, actin’s role in CME is known to be more pertinent at higher tension. In line with that, the loss of Tf number density on Cyto D treatment (**Fig. 5 C**) is understandably highest for the regions (midplane, higher tension) which already had an accumulation of coated pits. Next, we studied the impact of the tension profile in the average size of single pits. Regions at higher tension showed bigger Tf puncta (**Fig. 5 E**). The equivalent diameter of clathrin puncta also increased in central planes followed by a decrease at the top. The size of regions of colocalization of Tf with clathrin remained unaltered (**Fig. S9 J**).

**Figure 5:**
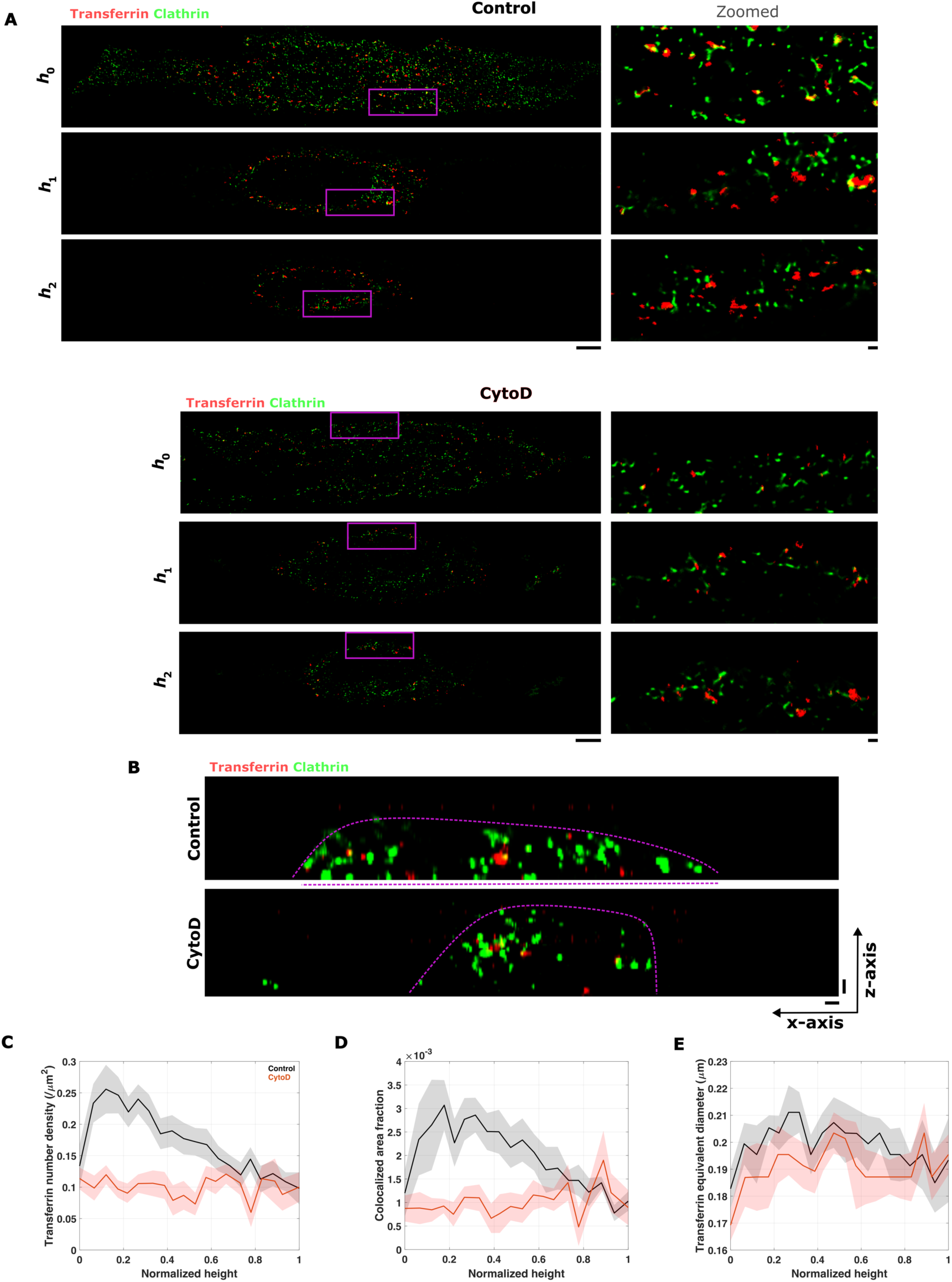
Apicobasal profile of endocytosis reveals the functional outcome of the actin and tension-gradient. A) Representative 3D STED z-sections of clathrin (immunolabeled with clathrin heavy chain Rabbit mAb and Abberior Star Red, represented in false color green) and Transferrin (labeled with Alexa Fluor 568, shown in false color red) at three different z-planes for both typical control and Cyto D-treated cells. Right panel: Zoomed in view of ROI outlined in magenta in the left panel. Scale bar (under main image) = 5 µm; scale bar (zoomed-in image) = 0.5 µm. B) Representative xz cross-sections with x being perpendicular the length of the line micropattern. Scale bar= 0.5 µm. C) Profile of No. of transferrin puncta detected per unit µm² for control (black) and Cyto D-treated cells (red), analyzed within 2.5 µm from the cell periphery, considered as the cortex, along the normalized height of the cell. D) Profile of total area of all transferrin puncta detected per unit area of the cortex for both control and Cyto D-treated cells, along the normalized height of the cell. E) Profile of colocalized area of transferrin and clathrin per unit area of the cortex, along the normalized height of the cell. N= 14 control cells and 11 Cyto D – treated cells.

This section thus captures the functional impact of the tension profile and also validates the loss of gradient of tension on Cyto D treatment – which was observed in OT-based measurements, but remained unclear from Flipper-TR lifetime measurements.

## Discussion

The mechanical state of the cell membrane is intricately regulated by the underlying actin cytoskeleton. A majority of past studies have treated cell membranes to be mechanically homogenous and reported effective tension to be enhanced by the actomyosin cytoskeleton.

In this paper, though the combination of three techniques, we investigated the different regions of the plasma membrane to understand if their mechanical state is uniform, noisy or has gradients. Combining IRM and OT-based tension measurements on the same imaging platform enabled us to make two different measurements on same cells. The comparison between IRM and OT yielded “tension” of similar magnitudes. While there exists a lack of one-to-one mapping between IRM and OT, a dependence of the tension ratio on tether height clearly suggested a height-dependent tension profile of the cell (**Fig. 1**). While an intact cortex is necessary for the tension gradient to be strong, the low spread area abolishes the gradient. Thus, the gradient seems relevant only for fairly well adhered cells (**Fig. 2B**).

We address the origin of this patterned tension by disrupting the cytoskeleton, and indeed bring out a new role of the cytoskeleton in tension homeostasis – that of reinforcement and maintenance of tension gradients. The average tether force drop per um is an indicator of the tension gradient. On Cyto D treatment, not only does the average value plummet, but it also becomes highly noisy. The difference remains statistically non-significant. Note that the cytoskeleton has been implicated already in the flow of tension. The enhanced heterogeneity in measured tension within cells points to poorer equilibration and hints to a slower flow of tension in line with other studies.

We employed Flipper-TR to get an overall view of tension throughout the cell to further validate the existence of the tension gradient. A recent study utilizing Flipper-TR has already reported the basal plane to be under lower tension than the midplane. However, in this work we not only go beyond basal to midplane comparison and map the complete z-profile using Flipper-TR, we also use OT-based dual tension measurements on same cells to assay the decrease in tension as height increases after the midplane. Note that the lifetime of Flipper-TR is also affected by lipid order (besides tension), and thus OT-based measurements validate the tension gradient and bring in better clarity to the mechanical state of the cell. However, Flipper-TR lifetime does not show any reduction in apical tension gradient on Cyto D treatment (**Fig. 4**) as observed in dual-tether measurements (**Fig. 2**). This effect could be similar to the higher change in lifetime found on hyper osmotic shock in comparison to hypo osmotic shock^28^.. The order-inducing effect of tension (that reduces lifetime on reducing tension) is usually opposed by the direct effect of reducing tension. The direct effect of reducing tension is the reducing of lipid-lipid distances, thus restricting rotational mobility and increasing lifetime of Flipper-TR. However, with the already existing state of low tension – since lipid-lipid distances cannot go down further, the direct effect of reducing tension may not affect the Flipper-TR lifetime. Hence, the impact of tension-induced order on the lifetime would be maximal. On the other hand, in low-tension Cyto D cells, we hypothesise that while order reduces (decreasing lifetime), the opposing effect of lifetime-increase by the direct effect on lipid-lipid distance doesn’t take place – resulting in a net higher decrease in lifetime per µm height increase. Here, it is important to note that we decided to go beyond biophysical techniques and draw inferences from the observed gradients in actin (**Fig. 3**), as well as perform experiments to assess gradients in functional processes such as endocytosis (**Fig. 5**).

To understand if the cytoskeleton contributes to the tension gradient, Cyto D was used to disrupt the acto-myosin network. Earlier studies have shown that pharmacologically reducing contractility by targeting myosin motors could either enhance or even reduce cell tension. The extent of cytoskeleton remodelling due to reduced contractility in particular cell lines is believed to vary and influence the results. To be able to detect subtle differences, in this work, cells were micropatterned on line-shaped structures so that the lateral cell surface displayed a clear cortical surface in a reproducible manner across cells and experiments. Dual tether OT-experiments further reduced the effect of cell-to-cell variability in obscuring the measurement of subtle tension gradients existing in cells.

To understand what in the cytoskeleton was responsible for the gradient, we imaged the overlap of myosin as well as ezrin with actin using super-resolution STED microscopy. With ∼ 40 nm resolution, STED could reveal the real overlaps of myosin fibers (length ∼ 100 nm) with actin, identifying the “contractile” elements of actin. Mapping the z-profile of contractility provided information similar to that from addition of drugs targeting myosin (usually done). The peak of colocalization was closely positioned to the peak in Flipper-TR lifetime (indicating high tension), followed by a clear decline validating the lower tension at apical planes and the role of contractility in it. Ezrin’s overlap with actin showed a non-uniform z-profile – however it did not capture the decreasing trend at the apical regions, underscoring contractility to be a major determinant of the tension profile than membrane-actin linkage.

Finally, the ambiguity of lack of loss of gradient of Flipper-TR lifetime (by Cyto D treatment) motivated the use of a functional assay to test the role of actin in creating the z-profile of tension. The z-profiles of clustered cargo and cargo-loaded CCPs matched that of acto-myosin colocalization. Using the cargo and clathrin together enabled us to identify the pits separately from the other structures (internalized vesicles, nascent clusters), besides showing a flattened profile on Cyto D validating the OT-based tension measurement data. Note that dynamic data could be used in future for validating the differences in the rate of endocytosis. However, our results drive us to comment on the presence of more invaginated coated pits at the central regions possibly contributing to the enhanced tension. For successful CME, actin is known not to be absolutely necessary but required at conditions of higher tension. This work shows that even within single cells, high tension regions having higher actin could form numerous pits. The bigger size^37^ could indicate their spending more time at the surface – which is in line with the belief that high tension disfavours endocytosis and clathrin structures are expected to be bigger^38^ and less curved than internalizing pits^39^. Previous work in our lab^32^ has shown that on Dynasore treatment (supresses pit to vesicle formation by pinching off), the number of CCPs increase, and the endocytic rate decreases – the tension being higher than normal conditions. Clearly, the state of the midplane closely resembles this kind of condition.

Besides the patterned tension and functional state, the data also highlight heterogeneities present in the cell. The heterogenous tension at the basal cell membrane is evident from the pixel-wise mapping of tension from IRM measurements (**Fig. 1 B**). A heterogeneity index (SD-Tension/mean-Tension) of 0.7 (using whole cell footprint) was measured at the basal plane (using IRM) while heterogeneity index of 0.25 (using 2 tethers per cell with average spacing of 3 µm) was measured at the lateral surface by tether-extraction.

In summary, we could demonstrate that adherent cells have stable patterned inhomogeneities in their apico-basal membrane tension profile that is dependent on substrate-attachment and acto-myosin contractility. We also reported that that such patterned mechanics is intertwined with the functional endocytic state of the cells. This work lays the foundation for further investigations on the impact of geometry of the patterns of the substrates in determining the tension patterns in z and study of the evolution of the tension profile across different kinds of cells in close-to physiological conditions so that the need for such patterned tension may be better understood.

## Material and Methods

### Resource

Resource:
| Resource availability REAGENT or RESOURCE | SOURCE | IDENTIFIER |
| --- | --- | --- |
| **Pharmacological Reagents** |  |  |
| Cytochalasin D | Sigma-Aldrich | Cat# C2618; LOT#84039 |
| Magnesium Chloride hexahydrate Cryst. Purified | Merck | Cat# 442615; CAS# 7791-18-6; |
| Dimethyl sulfoxide | Sigma-Aldrich | Cat# 276855; LOT# SHBC3339V; |
| HEPES | Sigma-Aldrich | Cat# H3375; LOT#SLBN0452V |
| Gelatin | Sigma-Aldrich | Cat# G2500; LOT# SLBX2973 |
| Glycine | Merck | Cat# 56-40-6 |
| TritonX-100 | Sigma-Aldrich | Cat# 101371902 |
| Alexa Flour 568 Phalloidin | Life Technologies | CAT#A12380; LOT#1600223 |
| Rabbit pAb to Ezrin (phospho T567) | Abcam | CAT#AB47293; LOT#GR324091-31 |
| Rabbit mAb to non-muscle Myosin IIA [EPR8965] Alexa Flour 488 | Abcam | CAT#ab204675; LOT#GR3280593-1 |
| Flipper-TR Bioprobe | Spirochrome | CAT#CY-SC020; LOT#057 |
| Transferrin from human serum, Alexa Flour 568 conjugate | Invitogen, Thermo Fisher Scientific | REF#T23365; LOT#2136786 |
| Clathrin heavy chain Rabbit mAb | Abclonal | CAT#A4943; LOT#4000001228 |
| Abberior STAR RED goat anti-rabbit IgG | Abberior | CAT#STRED-1002-100UG; LOT#30420PK-3 |
| Ultrapure Glycerol | Invitrogen, Thermo Fisher Scientific | REF#15514-011; LOT#19E1056865 |
| Mowiol 4-88 | ALDRICH Chemistry | CAT#81381; LOT#BCBK2317V |
| Tris Hydrochloride | SRL | CAT#99438; BATCH# 8677879 |
| DABCO 33-LV | ALDRICH Chemistry | CAT#290734; LOT#MKBK8680V |
| **Cell culture** |  |  |
| Cell line (HeLa) | ATCC | Cat# CCL-2; RRID# CVCL_0030 |
| Dulbecco's Modified Eagle's Medium (DMEM, High Glucose) | Gibco | Cat# 11965092 |
| Fetal Bovine Serum, certified, heat-inactivated, US | Gibco | Cat# 10082147 |
| Antibiotic-Antimycotic (100X) | Gibco | Cat# 15240062 |
| Trypsin-EDTA (0.25%), phenol red | Gibco | Cat# 25200072 |
| Paraformaldehyde | Sigma-Aldrich | Cat# P6148 |
| Phosphate Buffered Saline | Sigma-Aldrich | Cat# P3813 |
| Dulbecco's Modified Eagle's Medium (DMEM, Phenol Red free) | Gibco | Cat# 21063-029; LOT# 2239707 |
| **For calibration** |  |  |
| NIST Traceable Particle Size Standard, 60 μm | Bangs laboratories | Cat# L130806L; LOT# 11247 |
| **For optical trap bead** |  |  |
| 2.0 μm diameter | Invitogen, Thermo Fisher Scientific | REF#F8887; LOT#1971894 |
| **Software and algorithms** |  |  |
| ImageJ (FIJI) | NIH | https://fiji.sc/ |
| MATLAB 2020b | The Mathworks, Inc. | 40684590 |
| Labview 2014 | National Instruments | M61X44613 |

### Cell culture

All experiments utilized the HeLa cell line (CCL-2, ATCC) was used for all our experimental studies. Cells were cultured in Dulbecco’s Modified Essential Medium (DMEM, Gibco, Life Technologies, USA) supplemented with 10% fetal bovine serum (FBS, Gibco) and 1% Anti-Anti (Gibco) under conditions of 95% humidity, 5% CO_2_, and 37 °C. Cell were treated with 0.25% Trypsin–EDTA solution (Gibco) to detach them from the substrate for splitting and used for experiments typically after 16 – 18 hr of cell seeding..

### Pharmacological treatments

To prevent actin filament polymerization, cells were incubated in 5 μM Cytochalasin D (Cyto D, Sigma-Aldrich) in serum-free media for 40 minutes. The treatment was conducted inside the incubator at 37 °C. During the experiments, the Cyto D remained in the media.

### Immunostaining

To prepare for immunostaining, 4% paraformaldehyde (PFA, Sigma-Aldrich) was used. After 15 min incubation in PFA, cells were washed twice with 1X phosphate-buffered saline (PBS, Sigma-Aldrich), treated with 0.1 M glycine (Sigma-Aldrich) for 5 minutes and washed again with PBS. Subsequently, Triton-X was applied for 2 minutes, followed by another wash with PBS. For blocking, cells were kept in 3 ml of 0.2% Gelatin (Sigma-Aldrich) solution overnight at 4°C. The primary antibody treatment involved non-muscle Myosin II A raised in rabbit (Abcam) at a 1:400 dilution or p-ezrin antibody raised in rabbit (Abcam) (1:400 dilution) in Gelatin, and was left for 2 hr at room temperature. The Goat Anti-Rabbit IgG abberior STAR RED secondary antibody (abberior) was used at a 1:400 dilution in gelatin for 1 hr after washing with PBS. F-actin staining was performed using Alexa Fluor^TM^ 568 phalloidin (Thermo Fisher Scientific) with the secondary antibody in gelatin. Following the staining, the sample was washed three times in 1X PBS and then mounted on a clean coverslip with MOWIOL 4-88 (Sigma Aldrich) mounting media.

### IRM imaging and analysis

A Nikon Eclipse Ti-E motorized inverted microscope (Nikon, Japan) was used for IRM as well as optical-trap experiments. IRM^40^ imaging was performed using adjustable field and aperture diaphragms, a 100X Plan Apo (NA 1.4, oil immersion) objective on a 37 °C onstage incubator (Tokai Hit, Japan), s-CMOS camera (ORCA Flash 4.0, Hamamatsu, Japan), 100 W mercury arc lamp, an interference filter (546 ± 12 nm), and a 50–50 beam splitter. 2048 frames wre captured at 20 frames/s with an exposure time of 50 ms.

The intensity values from the Interference Reflection Microscopy (IRM) image series, which were aquired at a 50 ms exposure time and 20 frames per second, were converted to relative height from the base of the coverslip. This was done by calibrating with the IRM image of a known 60 µm diameter polystyrene bead (Polysciences) taken at different exposures. Subsequently, the power spectral density (PSD) of each pixel was calculated from the relative height time series using the covariance method in MATLAB.

The patches within a height range of 0 to 100 nm, with a specified tolerance limit, were identified as the first branch region (FBR) from the relative height time series. These FBR patches were selected for further analysis. To find out the mechanical parameters, the PSDs (obtained for each pixel and averaged over all pixels in an FBR) were fitted to 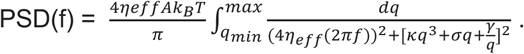

Here, A is the active temperature, *η*_*eff*_ is the effective cytoplasmic viscosity, *γ* is the confinement and *σ* is the membrane tension, which were used as fitting parameters. Here, the bending rigidity is kept constant to 15*k*_*B*_*T*. Any fit with R^2^< 0.99 were discarded and the region not not used for the analysis. Thus, from each cell, multiple FBR PSDs are fitted to find out the median tension from each FBR. The tension value of the cell is determined by taking the median value of the tension from multiple FBRs. While mapping tension in pixel-wise manner – which is solely performed for visualization, PSD of every pixel is fit to the above equation and all parameters including R^2^ of the fit are parallelly mapped for every pixel including regions under the nuclei which are never included in the FBR-wise tension analysis. For all rigorous comparisons only the FBR-wise tension measurements are used.

### Optical Trap experiments and analysis

In optical trap-based tether-pulling experiments, a 2 µm polystyrene bead was captured using a 1064 nm Laser (Coherent, Sweden) with a power of 1W at source, focused by a 100 X objective. The back-aperture of the objective was overfilled with an expanded collimated beam, with mirrors and a 50/50 beam splitter being used for beam manipulation. Imaging of the bead was done using a s-CMOS camera (ORCA Flash 4.0, Hamamatsu, Japan) while maintaining the sample at 37 °C using the onstage incubator (Tokai Hit, Japan). The trap stiffness (k) was calibrated using the equipartition approach, where 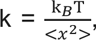 with x representing the displacement of the bead from the trap center, k_B_ being the Boltzmann constant, and T denoting the temperature in Kelvins. The bead was identified from captured images as an object (MATLAB, Fiji) for analysis, and its center was tracked over time. The bead’s displacement from the trap center and the spring constant of the trap were used to calculate the force (F = −*kx*). For each cell, a bead was first identified, trapped and its movement in trap recorded for stiffness calculation. The tether was pulled at a constant velocity of 0.5 µm/s up to 40 µm (LabVIEW, National Instruments, USA). The tether force was determined from the average bead position, with imaging conducted at 200 frames per second by defining a small region-of-interest selected, where both the bead and the cell are seen. The apparent membrane tension of the apical section of the cell was derived from the force using the Canham-Helfrich equation 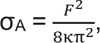 where κ signifies the bending rigidity (taken as 15 k_B_T), and σ_A_ represents the apparent membrane tension^10,24^.

### STED imaging and analysis

The Stimulated Emission Depletion (STED) Microscopy imaging was conducted using an inverted microscope (Abberior’s Facility Line STED). To ensure proper alignment of the STED and confocal point spread function (PSF), an auto-alignment sample containing uniform-sized multifluorescent beads was used. The 2D-STED imaging was done with a pixel size of 40 nm and the z-scan performed with a step size of 100 nm. A pulsed 775 nm laser was utilized for the depletion of red and far-red fluorescence. F-actin was stained with phalloidin tagged with Alexa Fluor 568, and the non-muscle myosin II A was tagged with STAR Red secondary antibody (Abberior).

For 3D-STED imaging, a 0.3 AU pinhole was used to reduce noise. The point spread function (PSF) for 3D-STED has a diameter of 70 nm along both the xy and z axes. This allows for clear visualization of spherical vesicles, as well as the distribution of transferrin and clathrin. In the 3D-STED imaging process, a step size of 50 nm and a height of 150 nm were used to collect the images.

To analyze the colocalization of F-actin and non-muscle myosin II A at the cortex, STED images were taken and then the object was detected by creating binary images for both images. To create the binary images for analysis, a Gaussian blur operation was first performed, followed by subtracting it from the original image to reduce noise. The noise-reduced image was then converted to values between 0 and 1, and proper thresholding was performed to form a binary image which subsequently used for object detection. For analysing the colocalization at the cortex, a region of about 400 nm (10 pixels) was considered within the cortex region.

To determine the cortical region of interest, the cell was roughly selected on the actin image to exclude objects away from the cortex. Then, the outer rim of the cell was detected from the selected region of the actin’s binary image. A binary image was created from the outer rim, which was then eroded by 10 pixels. The original outer rim binary image was then subtracted from it to find the exact cortical region of interest.

For colocalization, the binary image was selected from the finally determined cortical region’s region of interest and then simply added to find out the regions that show a value of 2. The fraction of actin colocalization to myosin was found using the formula: Σ(Pixels colocalized (value = 2))/Σ(Total actin pixels (value = 1 and 2)) at the cortical region of the cell. The colocalized actin intensity represents the total fluorescence of the actin at the colocalized region. Furthermore, the fraction of colocalization multiplied by the total actin intensity at the cortex was also calculated for further analysis.

This analysis was performed for the whole cell with a 100 nm change in step along the height of the cell.

### Fluorescence Lifetime imaging and analysis

For fluorescence lifetime imaging of Flipper-TR, cells were treated with 1 µM of Flipper-TR for 30 min in M1 medium and used for imaging. The Abberior facility line built around the Olympus IX83 equipped with time correlated single photon counting (TCSPC module, PicoQuant). A 60x oil objective (1.42 NA) was used for the imaging keeping x, y pixel size as 200 nm and Δ*z* of z-scan as 400 nm. Typically > 50 photons were acquired for pixels inside the cell.

For analysis, the lifetime provided by the phasor analysis for every pixel was used for all pixels with photon count greater than 20. The cellular periphery (even at the basal membrane) was manual tracked and the line ROI used to obtain the median lifetime per image plane.

### Endocytosis assay and analysis

For endocytosis uptake assay, 10 μg/ml Transferrin Alexa fluor 568 (Tf) was incubated for 5 min at 37°C. For stopping endocytosis, HEPES based buffer was used at ice cold temperature. Imaging was performed using the 3D STED and analysis was done by standard local threshold-based object detection as described in the STED analysis section. The peripheral ROI thickness in this case was taken to be 2.5 µm since the interest was to analyze both surface clusters and cortical endosomes.

### Statistical analysis

Every experiment was repeated three times involving multiple cells. A Mann–Whitney U test was performed for statistical significance testing (ns denotes p > 0.05, * denotes p < 0.05, ** denotes p < 0.001).

## Acknowledgement

B.S. acknowledges support from Wellcome Trust/DBT India Alliance fellowship (grant number IA/I/13/1/500885), SERB (grant number SERB_CRG_2336) and CEFIPRA (grant number 6303-1). T.G. thanks CEFIPRA and IISERK for providing his fellowship. A.M. thanks IISERK for providing her fellowship. The authors are also thankful to the Builder Imaging facility (BT/INF/22/SP45383/2022) for super-resolution STED imaging.

## Author Contributions

B.S. and A. B. designed research, acquired funding; T.G., A.M. and A.K. set up methodology; T.G. and A.M. performed research and analysed data; T.G. also performed STED/Lifetime experiments and analysis; and T.G., A.M., A.B. and B.S. wrote the paper; everyone edited the paper.

## Competing Interest Statement

The authors declare no competing interest.

## Notes

### Competing Interest Statement

The authors have declared no competing interest.

