## Supplementary Material for "Actin contractility and endocytosis create apico-basal tension gradient in HeLa cells"

### Suuplementary Text

**LIST OF FIGURES:**

**Figure S1:** Analysis procedure in MATLAB for detecting centroids of beads used in force measurements.

**Figure S2:** Dependence of the standard deviation of height fluctuations in IRM with the optical trap tension measured for single cells.

**Figure S3:** Maps of additional fitting parameters (pixel-wise map) and FBR-wise tension tension map (Fig. 1(B)).

**Figure S4:** Comparison of the spread area of Lin2 micropattern cells compared to Intermediate micropattern cells.

**Figure S5:** Effect of Cyto D on Flipper-TR Lifetime and its intracellular heterogeneity.

**Figure S6:** Distribution of transferrin along the height of the cell using a 16-color lookup table (LUT).

**Figure S7:** Procedure for Analyzing the Cortex Region in MATLAB.

**Figure S8:** Parameters analyzed after processing the Transferrin-Clathrin images to binary.

**Figure S9:** Analysis of transferrin-clathrin distribution along the cell height for control and Cyto D-treated cells.

**Table S1:** List of statistical parameters of all figures.

**Table S2:** Parameters used for analyzing fluorescence-labelled objects detected.

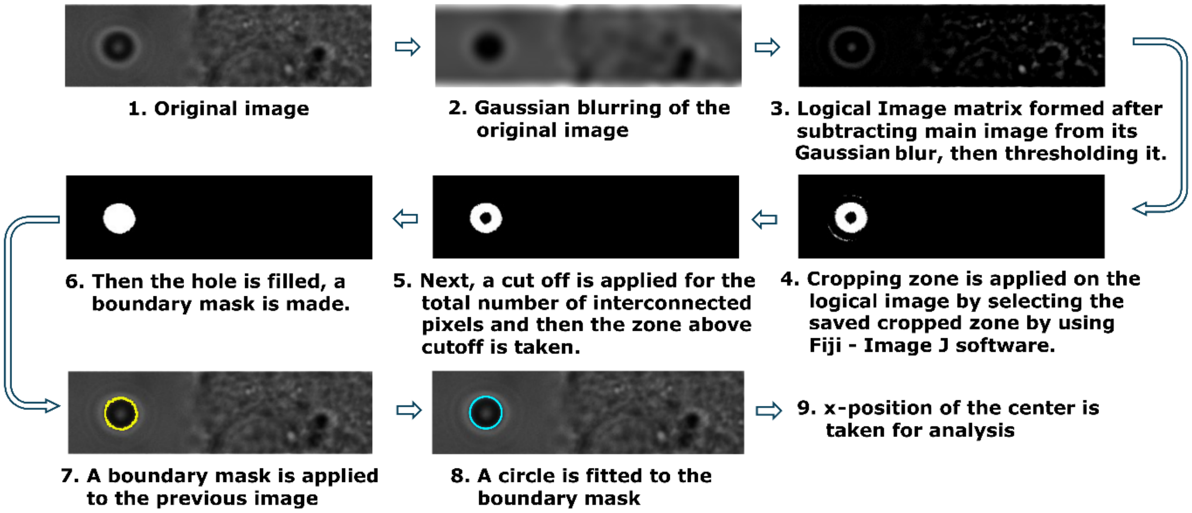

*Figure S1:* ***Analysis procedure in MATLAB for detecting centroids of beads used in force measurements.***

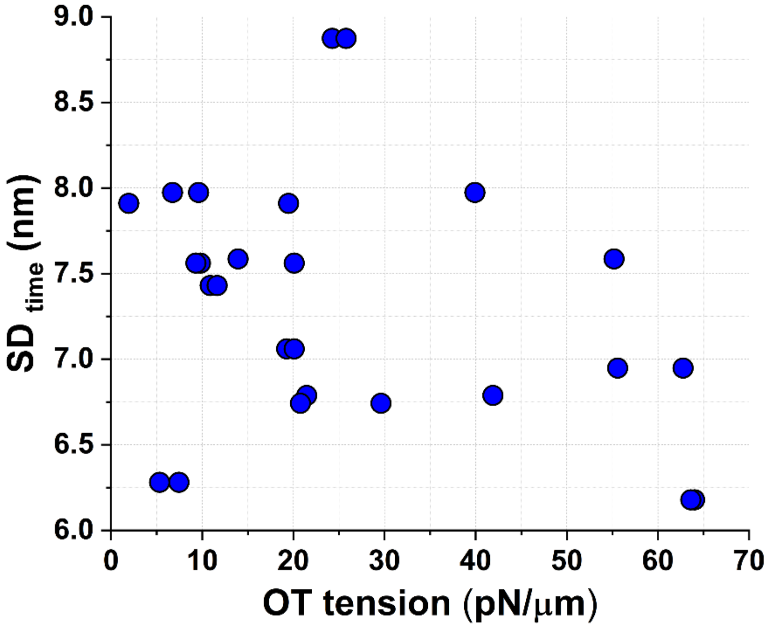

*Figure S2:* ***Dependence of the standard deviation of height fluctuations in IRM with the optical trap tension measured for single cells.*** *N= 28 cells.*

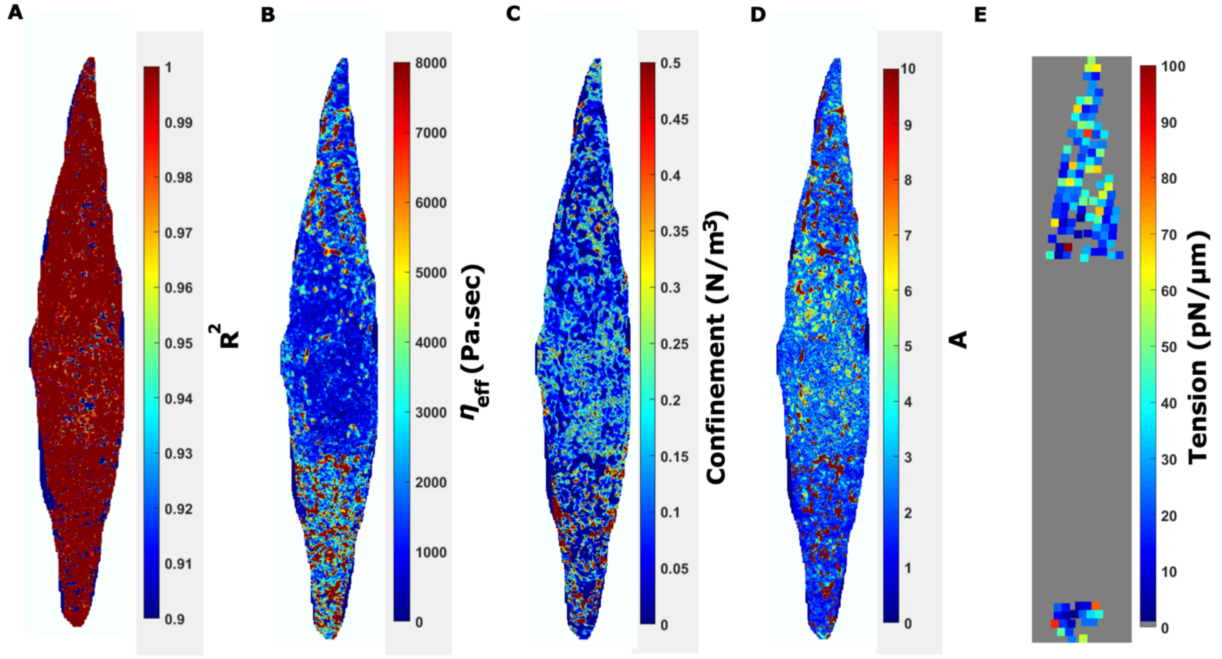

*Figure S3:* ***Maps of additional fitting parameters (pixel-wise map) and FBR-wise tension tension map (Fig. 1(B)).*** *(A) This represents the $R^2$ values of fitting of PSDs at each pixel – for visualization purposes – to obtain the tension (B) Map of fitting parameter - effective cytoplasmic viscosity. (C) Map of fitting parameter - confinement. (D) Map of fitting parameter – active temperature. (E) Map of tension obtained from fitting PSDs averaged over individual FBRs. The section missing at the centre of the cell corresponds to the region under the nuclei which was not analysed and regions where the fitting was not proper ($R^2$<0.999).*

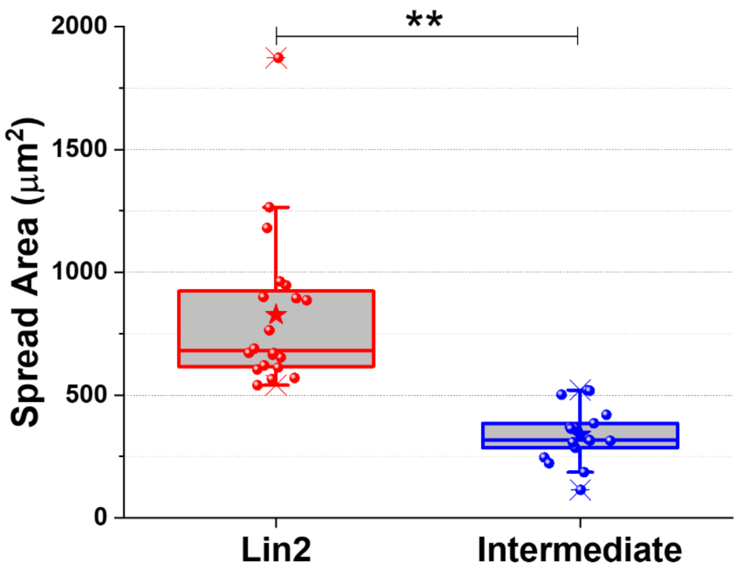

*Figure S4:* ***Comparison of the spread area of Lin2 micropattern cells compared to Intermediate micropattern cells.*** *The significant difference between the two data sets was found to be 1.257 x $10^{-10}$. Cells used for Lin2 micropattern and intermediate micropattern is 20 and 17 respectively. Plot representing that the intermediate micropattern has low spread area as the normal spread area and low spread area's fractional drop in tension is different than normal fig. 2(F).*

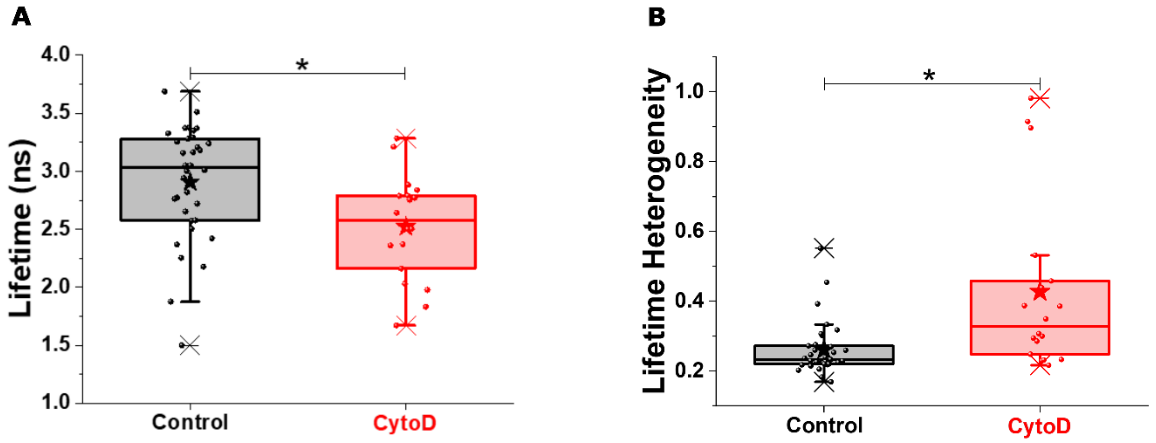

*Figure S5:* ***Effect of Cyto D on Flipper-TR Lifetime and its intracellular heterogeneity.*** *Figure illustrating the (A) Flipper-TR lifetime observed at the base of the cell and (B) the heterogeneity in lifetime (standard deviation of lifetime divided by mean lifetime) at the base of the cell. A total of 34 cells were analyzed for the control group, while 18 cells were examined in the Cyto D-treated group.*

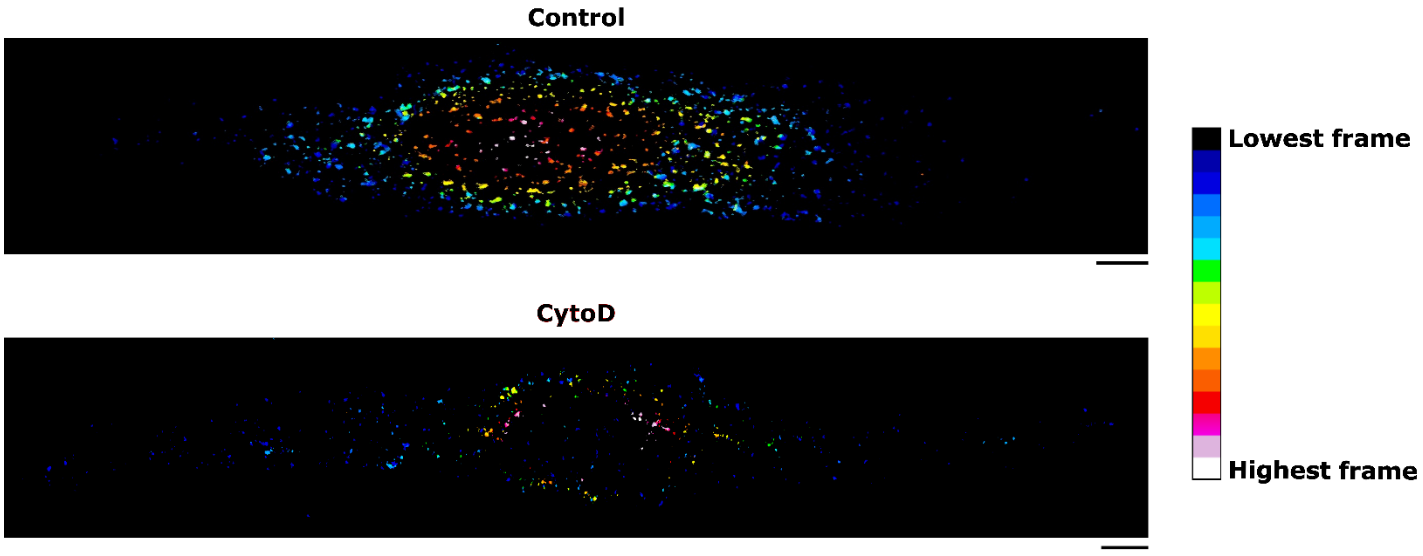

*Figure S6:* ***Distribution of transferrin along the height of the cell using a 16-color lookup table (LUT).*** *The LUT is applied vertically to visualize the number and size of transferrin present throughout the cell's height, allowing for observation without the need for further analysis. The scalebar under the image represents 5µm.*

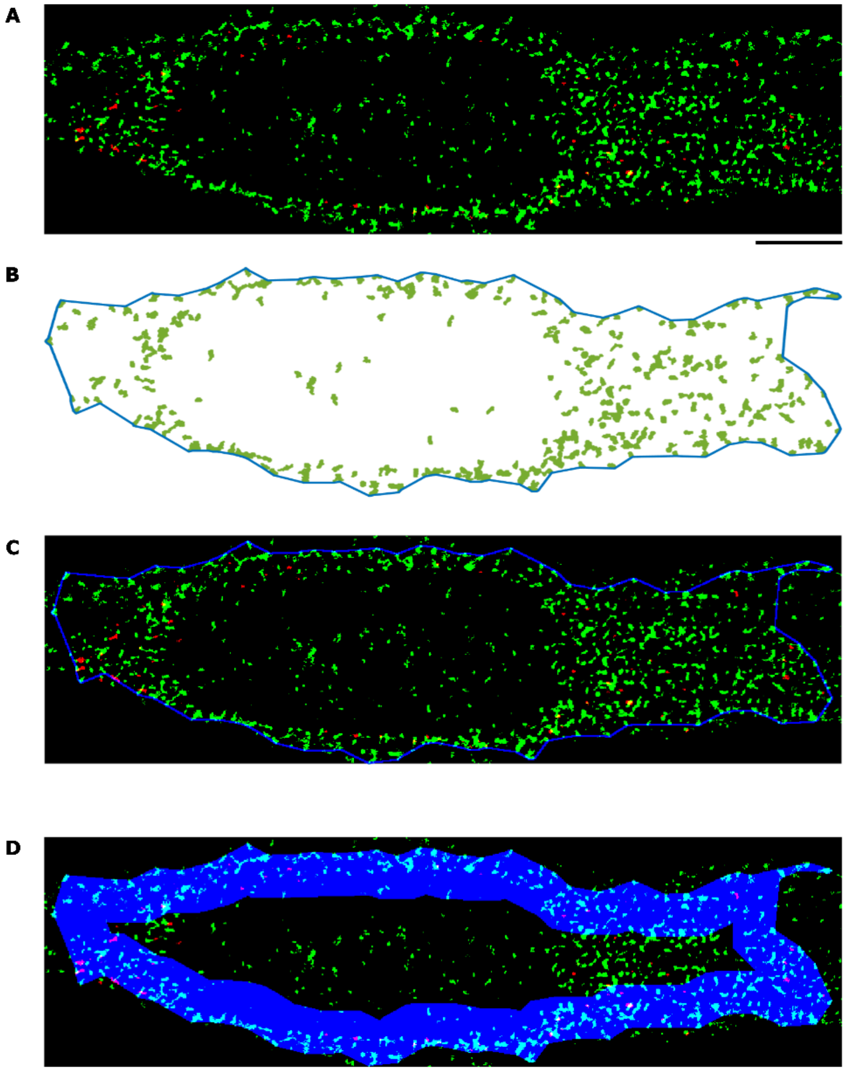

*Figure S7:* ***Procedure for Analyzing the Cortex Region in MATLAB.*** *(A) In the image, green indicates the immunolabelled clathrin, while red represents the transferrin. The convoluted 3D-STED image is presented here and used in analysis. Scalebar represents 5 µm (B) The precise edge of the cell is identified from the binary image of Clathrin, which is shown here as a blue line outlining the cell. The region of interest (ROI) at the edge of the cell is detected using a MATLAB code snippet that incorporates a shrink factor and a cutoff value for the number of interconnected pixels. The shrink factor can vary between 0 and 1, with 0 indicating the lowest level of shrinkage and 1 representing the maximum level. For optimal ROI edge detection, typical values for the cutoff and shrink factor are utilized. (C) The boundary is applied to the main image to verify its position on the cell's periphery. (D) Initially, a mask is created of the boundary, which is then eroded by 2.5-microns. This eroded area is subtracted from the original mask, resulting in a 2.5-micron thick area at the periphery, depicted in blue.*

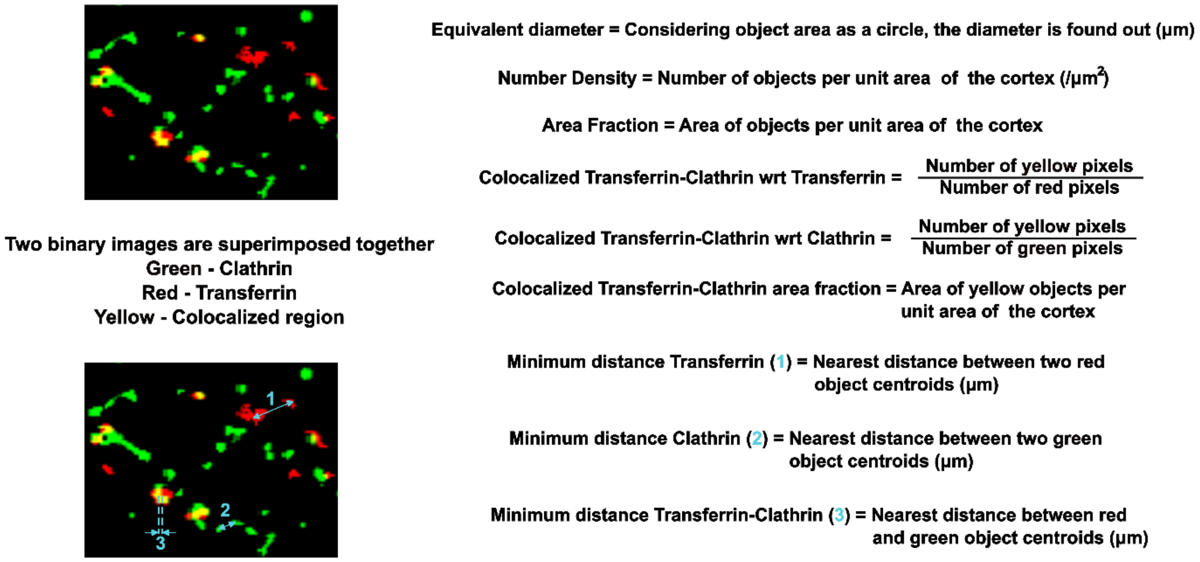

*Figure S8:* ***Parameters analyzed after processing the Transferrin-Clathrin images to binary.*** *After the calculation of the parameters the median values across the height of the cortex are used for plotting.*

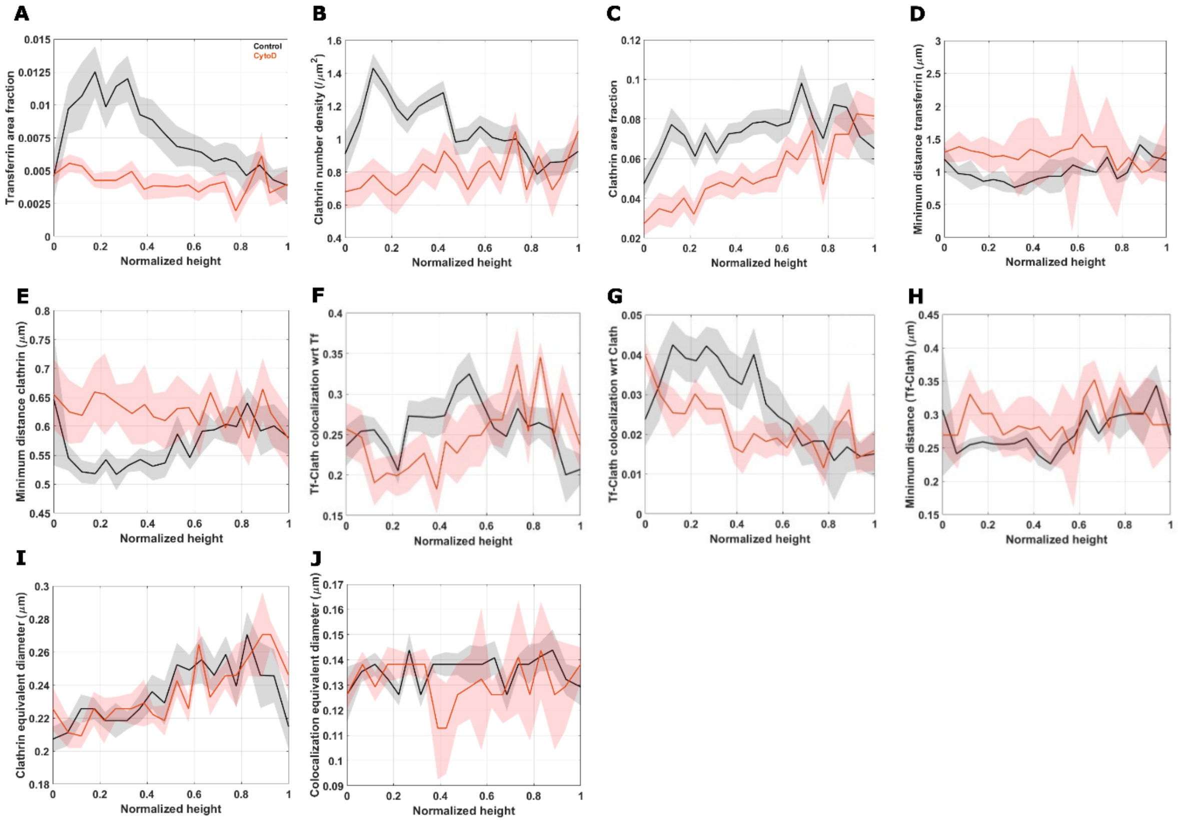

*Figure S9:* ***Analysis of transferrin-clathrin distribution along the cell height for control and Cyto D-treated cells.*** *(A) In this context, for all the plots, black indicates control cells, while red represents cells treated with Cyto D. The plot illustrates the variation in transferrin area fraction along the cells' height. (B) The plot illustrates the density of clathrin objects per unit area of the cortex along the height of the cells. (C) The plot illustrates the area of clathrin objects per unit cortical area defined along the cells' height. (D) The plot shows the minimum distance between transferrin objects along the height of the cell. (E) The plot shows the minimum distance between clathrin objects along the height of the cell. (F) The plot illustrates the ratio of transferrin to the area of colocalized transferrin and clathrin. (G) The plot illustrates the ratio of clathrin to the area of colocalized transferrin and clathrin. (H) The plot shows the minimum distance between transferrin and clathrin puncta along the height of the cell. (I) Clathrin's equivalent diameter is plotted along the height of the cells. The equivalent diameter is determined from the area of the object, assuming the objects are spherical in nature. (J) The equivalent diameter of the colocalized area objects is plotted along the height of the cell. All the plots were done for 14 control cells and 11 Cyto D – treated cells.*

| Table S1 |
|---|
| **List of statistical parameters of figures.** |

| **Figure 1B** | | | | |
|---|---|---|---|---|
| | | **Forces (pN)** | | |
| | | **Mean** | **Median** | **STD** |
| **Phases** | **Time period (sec)** | | | |
| 1 | 0-0.5 | 0 | 0 | 0.6076 |
| 2 | 25-35 | -11.5896 | -11.5841 | 0.9237 |
| 4(1) | 75-90 | 20.6354 | 20.5854 | 1.0254 |
| 4(2) | 100-125 | 9.4202 | 9.4013 | 0.8251 |
| 5 | 220-222.5 | 0.034 | -0.0454 | 1.0671 |

| **Figure 1D** | | | | | | | |
|---|---|---|---|---|---|---|---|
| **Parameters** | **Conditions** | **N** | **Mean** | **Median** | **SD** | **SEM** | **p-values (wrt 1)** |
| Ratio (Ot-Ten/IRM-Ten)] | Control | 13 | 0.7117 | 0.5849 | 0.4465 | 0.1238 | 0.0137 |

| **Figure 2C** | | |
|---|---|---|
| | **Control** | **Cyto D** |
| **Pearson Correlation coefficient** | -0.50966 | -0.1763 |
| **p-value** | 0.00926 | 0.38895 |

| **Figure 2D** | | | | | | | | |
|---|---|---|---|---|---|---|---|---|
| **Parameters** | **Conditions** | **N** | **Mean** | **Median** | **SD** | **SEM** | **p-values (wrt 1)** | **p-values between them** |
| F(H1)/F(H2) | Control | 37 | 1.48986 | 1.20188 | 0.96716 | 0.159 | 8.08E-09 | 0.04002 |
| | Cyto D | 25 | 1.53929 | 1.15702 | 1.46825 | 0.29365 | 0.55054 | 0.04002 |

| **Figure 2E** | | | | | | | | |
|---|---|---|---|---|---|---|---|---|
| **Parameters** | **Conditions** | **N** | **Mean** | **Median** | **SD** | **SEM** | **p-values (wrt 0)** | **p-values between them** |
| Fraction drop of tension per µm height increase | Control | 53 | 0.3378 | 0.1308 | 0.7936 | 0.1090 | 2.13e-07 | 0.4274 |
| | Cyto D | 27 | 1.1523 | 0.0613 | 3.2593 | 0.6273 | 0.81 | 0.4274 |

| **Figure 2F** |
|---|

| Parameters | Conditions | N | Mean | Median | SD | SEM | p-values (wrt 0) | p-values between them |
|---|---|---|---|---|---|---|---|---|
| Fraction drop of tension per µm height increase | Control | 53 | 0.3378 | 0.1308 | 0.7936 | 0.1090 | 2.13e-07 | 0.0163 |
| | Low Spread Area | 13 | -2.5693 | -0.006 | 9.2828 | 2.5746 | 0.7423 | 0.0163 |

| Figure 3E | | |
|---|---|---|
| Using data sets between normalized heights 0.6 and 0.9 | | |
| | Mean cortical actin intensity (A.U.) | |
| | Control | Cyto D |
| Pearson's Correlation coefficient | -0.4319 | -0.1301 |
| p-value | 7.07E-05 | 0.326 |

| Figure 3F | | |
|---|---|---|
| Using data sets between normalized heights 0.6 and 0.9 | | |
| | Fraction of actin colocalized to myosin vs. normalized height | |
| | Control | Cyto D |
| Pearson's Correlation coefficient | -0.3398 | -0.1038 |
| p-value | 0.0022 | 0.438 |
| | Fraction of actin colocalized to ezrin vs. normalized height | |
| | Control | Cyto D |
| Pearson's Correlation coefficient | 0.1954 | 0.0039 |
| p-value | 0.0041 | 0.9603 |

| Figure 4D | | | | | | | | |
|---|---|---|---|---|---|---|---|---|
| **Parameters** | **Conditions** | **N** | **Mean** | **Median** | **SD** | **SEM** | **p-values (wrt 1)** | **p-values between them** |
| **Lifetime ratio** | **Control (11.2 µm wrt 8.8 µm)** | 20 | 0.97959 | 0.97572 | 0.0332 | 0.00742 | 5.50E-05 | 0.01667 |
| | **Cyto D (10 µm wrt 7.2 µm )** | 12 | 0.92581 | 0.94998 | 0.07176 | 0.02071 | 2.42E-04 | 0.01667 |

| Figure 5C |
|---|
| Transferrin number density (Using data sets between normalized heights 0.6 and 0.9) |

| | Control | Cyto D |
|---|---|---|
| **Pearson's Correlation coefficient** | -0.2079 | 0.1128 |
| **p-value** | 0.0715 | 0.4124 |

| **Figure 5D** | | |
|---|---|---|
| **Colocalized area fraction (Using data sets between normalized heights 0.6 and 0.9)** | | |
| | **Control** | **Cyto D** |
| **Pearson's Correlation coefficient** | -0.2551 | 0.2614 |
| **p-value** | 0.0262 | 0.0587 |

| **Figure 5E** | | |
|---|---|---|
| **Transferrin equivalent diameter (Using data sets between normalized heights 0.6 and 0.9)** | | |
| | **Control** | **Cyto D** |
| **Pearson's Correlation coefficient** | -0.0742 | -0.1722 |
| **p-value** | 0.5239 | 0.2175 |

| **Figure S4** | | | | | | | |
|---|---|---|---|---|---|---|---|
| **Parameters** | **Conditions** | **N** | **Mean** | **Median** | **SD** | **SEM** | **p-values between them** |
| **Spread Area ($\mu m^2$)** | **Lin2 micropattern** | 20 | 826.8024 | 680.1575 | 319.5831 | 71.46094 | 2.40E-07 |
| | **Intermediate micropattern** | 17 | 338.5844 | 317.758 | 112.9356 | 27.39091 | 2.40E-07 |

| **Figure S5 (A)** | | | | | | | |
|---|---|---|---|---|---|---|---|
| **Parameters** | **Conditions** | **N** | **Mean** | **Median** | **SD** | **SEM** | **p-values between them** |
| **Lifetime at base (sec)** | **Control** | 34 | 2.91E-09 | 3.03E-09 | 4.90E-10 | 8.41E-11 | 6.88E-03 |
| | **Cyto D** | 18 | 2.52E-09 | 2.57E-09 | 4.52E-10 | 1.07E-10 | 6.88E-03 |

| **Figure S5 (B)** | | | | | | | |
|---|---|---|---|---|---|---|---|
| **Parameters** | **Conditions** | **N** | **Mean** | **Median** | **SD** | **SEM** | **p-values between them** |

| | | | | | | | |
|---|---|---|---|---|---|---|---|
| **Lifetime heterogeneity at base** | **Control** | 34 | 2.61E-01 | 2.32E-01 | 7.65E-02 | 1.31E-02 | 1.00E-03 |
| | **Cyto D** | 18 | 0.42683 | 0.32785 | 0.24817 | 0.0585 | 1.00E-03 |

| **Figure S9 (A)** | | |
|---|---|---|
| **Transferrin area fraction (Using data sets between normalized heights 0.6 and 0.9)** | | |
| | **Control** | **Cyto D** |
| **Pearson's Correlation coefficient** | -0.2141 | 0.2540 |
| **p-value** | 0.0633 | 0.0665 |

| **Figure S9 (B)** | | |
|---|---|---|
| **Clathrin number density (Using data sets between normalized heights 0.6 and 0.9)** | | |
| | **Control** | **Cyto D** |
| **Pearson's Correlation coefficient** | -0.2141 | 0.1005 |
| **p-value** | 0.0634 | 0.4740 |

| **Figure S9 (C)** | | |
|---|---|---|
| **Clathrin area fraction (Using data sets between normalized heights 0.6 and 0.9)** | | |
| | **Control** | **Cyto D** |
| **Pearson's Correlation coefficient** | 0.0683 | 0.2554 |
| **p-value** | 0.5578 | 0.0649 |

| **Figure S9 (D)** | | |
|---|---|---|
| **Minimum distance transferrin (Using data sets between normalized heights 0.6 and 0.9)** | | |
| | **Control** | **Cyto D** |
| **Pearson's Correlation coefficient** | 0.2021 | -0.2729 |
| **p-value** | 0.0800 | 0.0480 |

| **Figure S9 (E)** | | |
|---|---|---|
| **Minimum distance clathrin (Using data sets between normalized heights 0.6 and 0.9)** | | |
| | **Control** | **Cyto D** |
| **Pearson's Correlation coefficient** | 0.1855 | -0.0646 |
| **p-value** | 0.1087 | 0.6456 |

| **Figure S9(F)** |
|---|

| Tf-Clath colocalization wrt Tf (Using data sets between normalized heights 0.6 and 0.9) | | |
|---|---|---|
| | Control | Cyto D |
| Pearson's Correlation coefficient | 0.0363 | 0.0637 |
| p-value | 0.7553 | 0.6506 |

| Figure S9 (G) | | |
|---|---|---|
| Tf-Clath colocalization wrt Clath (Using data sets between normalized heights 0.6 and 0.9) | | |
| | Control | Cyto D |
| Pearson's Correlation coefficient | -0.0856 | 0.0962 |
| p-value | 0.4624 | 0.4931 |

| Figure S9 (H) | | |
|---|---|---|
| Minimum distance Tf-Clath (Using data sets between normalized heights 0.6 and 0.9) | | |
| | Control | Cyto D |
| Pearson's Correlation coefficient | 0.1687 | -0.1666 |
| p-value | 0.1451 | 0.2331 |

| Figure S9 (I) | | |
|---|---|---|
| Clathrin equivalent diameter (Using data sets between normalized heights 0.6 and 0.9) | | |
| | Control | Cyto D |
| Pearson's Correlation coefficient | 0.1582 | 0.1929 |
| p-value | 0.1724 | 0.1665 |

| Figure S9 (J) | | |
|---|---|---|
| Colocalization equivalent diameter (Using data sets between normalized heights 0.6 and 0.9) | | |
| | Control | Cyto D |
| Pearson's Correlation coefficient | 0.1012 | -0.1783 |
| p-value | 0.3844 | 0.1691 |

| Table S2 |
|---|
| **Parameters used for analyzing fluorescence-labelled objects detected.** |

| S.No. | Parameter (of objects) | Definition |
|---|---|---|
| 1. | **Equivalent diameter** | $= 2\sqrt{\frac{A}{\pi}}$ where A is the area per object or the total number of pixels in the object multiplied by the area of each pixel). |
| 2. | **Number Density** | $= \frac{N}{A_c}$ where N is the number of objects detected from the image. "$A_c$" denotes the Area of the ROI for a particular z in which the objects are detected. In this case, the area of the cortical ROI is taken. |
| 3. | **Area Fraction** | $= \frac{\sum_i A_i}{A_c}$ where $A_i$ represents the area of each object while $A_c$ represents the area of the cortical ROI. |
| 4. | **Minimum distance** | Centroid-to-centroid distance between nearest neighbours of relevant labels. |
| 5. | **Colocalization are fraction** | $= \frac{A_Y}{A_c}$ where $A_Y$ represents the total area of "yellow" or colocalized pixels and $A_c$ represents the area of the cortical ROI. |
| 6. | **Colocalization wrt label** | $= \frac{A_Y}{A_L}$ where $A_Y$ represents the total area of "yellow" or colocalized pixels and $A_L$ represents the total area covered by objects detected of a particular label. |
| **Method of averaging** | | Median for parameters measured for all objects detected in a particular z for each cell is first collated with the z-values. Subsequently the median is obtained particular z-bands from these cell-wise averages and plotted with error bar as SEM calculated over the cells. |
